# Clinical prediction of wound re-epithelisation outcomes in non-severe burn injury using the plasma lipidome

**DOI:** 10.1101/2023.07.28.550938

**Authors:** Monique J. Ryan, Edward Raby, Reika Masuda, Samantha Lodge, Philipp Nitschke, Garth L. Maker, Julien Wist, Mark W. Fear, Elaine Holmes, Jeremy K. Nicholson, Nicola Gray, Luke Whiley, Fiona M. Wood

## Abstract

Whilst wound repair in severe burns has received substantial research attention, non-severe burns (<20% total body surface area) remain relatively understudied, despite causing considerable physiological impact and constituting most of the hospital admissions for burns. Early prediction of healing outcomes would decrease financial and patient burden, and aid in preventing long-term complications from poor wound healing. Lipids have been implicated in inflammation and tissue repair and may play essential roles in burn wound healing. In this study, plasma samples were collected from 20 non-severe burn patients over 6 weeks from admission, including surgery, and analysed by liquid chromatography-tandem mass spectrometry and nuclear magnetic resonance spectroscopy to detect 850 lipids and 112 lipoproteins. Orthogonal projections to latent structures-discriminant analysis was performed to identify changes associated with re-epithelialisation and delayed re-epithelisation.

We demonstrated that the lipid and lipoprotein profiles at admission could predict re-epithelisation outcomes at 2 weeks post-surgery, and that these discriminatory profiles were maintained up to 6 weeks post-burn. Inflammatory markers GlycB and C-reactive protein indicated divergent systemic responses to the burn injury at admission. Triacylglycerols, diacylglycerols and low-density lipoprotein subfractions were associated with delayed wound closure (p-value <0.02, Cliff’s delta >0.7), whilst high-density lipoprotein subfractions, phosphatidylinositols, phosphatidylcholines, and phosphatidylserines were associated with re-epithelisation at 2 weeks post-surgery (p-value <0.01, Cliff’s delta <-0.7). Further model validation will potentially lead to personalised intervention strategies to reduce the risk of chronic complications post-burn injury.

## Introduction

Burn injuries represent a significant global health burden and pose considerable challenges to clinicians in achieving successful wound healing[1,2]. While severe burns receive considerable research into improvement of clinical management[3–5], the majority of burn injuries requiring hospitalisation are clinically classed as “non-severe” - defined as <20% total burn surface area (TBSA)[6]. Despite a non-severe classification, these injuries can still result in significant physiological damage and can compromise skin barrier functionality[7]. Due to the heterogeneity of non-severe burn injury, prediction of wound outcomes is challenging, with clinical classification and subsequent management reliant on visual wound observation, which has been reported to only be 60-75% accurate[2,8,9].

Due to the complex nature of wound healing, any imbalances or dysregulation in the associated pathways can lead to impaired wound healing and chronicity[11]. Several issues can contribute to poor wound healing, including but not limited to, increased oxidative stress, biofilm formation, cytokine, chemokine and growth factor imbalances, leaky blood vessels and excessive inflammation[11–13]. Impaired wound healing can result in short- and long-term health complications. In both severe and non-severe burns, a significant issue with delayed wound healing is the risk of graft failure[14], increased susceptibility to infection[15] and scar formation[16]. Scarring, particularly hypertrophic scars, can leave patients with long-term mobility issues, and chronic itch, discomfort and/or painful sensations[16]. Even without visible scars, an estimated 70% of burns patients will experience dysesthesia during their recovery, which can result in chronic pain, hyper- or hypo-sensitivity and temperature fluctuations[18]. Cumulatively, these can negatively impact physical and mental health, increasing the risk of depression and anxiety[17] and affecting their quality of life.

In recent years, there has been an emergence of biomarkers that could predict patient mortality and outcomes in severe burns. Biomarkers such as bilirubin[19], creatinine[19], and C-reactive protein (CRP)[20] have been monitored longitudinally and show promise as prognostic markers of mortality. However, predictive research has mostly focused on mortality[19–21] and sepsis[22], due to the life-threatening nature of severe burns, with minimal research on the wound healing process itself[23]. Targeting wound healing is critical to the post-burn recovery process, as it can prevent infections and prolonged inflammation which are common causes of morbidity and mortality in burn patients[4,23,24]. Predicting wound healing outcomes in the early stages of a burn provides a potential opportunity for improving patient outcomes. Early prediction can lead to increased effectiveness of time-critical treatments, including wound debridement, that when performed late has been shown to delay wound closure and is associated with increased hospitalisation duration and infections[25,26]. Additionally, early prediction can stratify patients into risk groups for wound healing outcomes, which can guide clinicians into different treatment regimens for optimising patient recoveries, reducing health burden and financial cost[27]. To date, minimal research has been conducted on predicting wound healing outcomes in non-severe burns[8], despite accounting for up to 90% of the patient population in developed countries[28].

To research complex conditions such as wound healing, studies have moved away from traditional single biomarkers to panels of biomarkers or metabolic phenotypes, as the metabolome is directly influenced by the physiological state and vice versa[29]. Novel analytical techniques such as liquid chromatography-mass spectrometry (LC-MS) and nuclear magnetic resonance (NMR) spectroscopy, are able to analyse small molecules within the plasma or serum metabolome, shedding light on perturbed systemic metabolism that may be associated with disease[30–33]. These techniques have emerged as powerful tools for analysing metabolic phenotypes and identifying metabolic signatures that can predict patient outcomes in the early stages of disease, including cardiovascular disease[29,34,35], diabetes[29,36], infection[31,37,38] and cancer[39,40]. Recovery trajectories post-infection can also be predicted from baseline samples with high accuracy and precision [38] Lipid and lipoprotein metabolism has been associated with numerous inflammatory mechanisms and previous research into non-severe burns has identified dysregulation in these signatures up to three years post-burn injury[41]. No other studies have investigated lipid and lipoprotein profiles longitudinally in non-severe burns to assess its ability to predict patient outcomes, even though the potential has been identified with respect to severe burns[42,43] and other inflammatory conditions[44].

We have previously demonstrated that lipid and lipoprotein metabolism measured by liquid chromatography-tandem mass spectrometry (LC-QQQ-MS) and NMR spectroscopy, respectively, in this non-severe burn cohort was perturbed up to six weeks post-injury when compared to non-burn controls[45]. The study suggested that patients were experiencing persistent inflammation that could impact long-term recovery[11]. As lipid and lipoprotein differences were apparent at burn admission, we investigated if these signatures could predict the re-epithelisation of non-severe burn wounds at two weeks post-surgery, a common wound healing parameter assessed by clinicians when monitoring wound recovery. By employing the same analytical techniques, this study aimed to identify whether monitoring lipid and lipoprotein profile trajectories could effectively predict wound healing outcomes in the early stages of non-severe burn injury and potentially be translated clinically to identify patients at risk of poor wound healing.

## Materials and methods

### Reagents and chemicals

LC-MS grade water was produced from a Milli-Q® IQ 7000 (Merck Millipore) and Optima LC-MS grade organic solvents were purchased from Thermo Fisher Scientific (Malaga, WA, Australia). Phosphate buffer was acquired from Bruker (Bruker BioSpin, Billerica, Massachusetts, USA). Isotopically labelled lipid standards were obtained from the Lipidyzer^TM^ Internal Standards Kit from SCIEX (MA, USA), and SPLASH LipidoMIX^TM^ internal standards, lysoPG 17:1 and lysoPS 17:1 were purchased from Avanti Polar Lipids (Sigma-Aldrich, North Ryde, NSW, Australia).

### Study design and biological samples

Non-severe burn plasma samples were acquired from the placebo arm of a randomised placebo-controlled trial of Celecoxib for Acute Burn Inflammation and Fever (CABIN Fever Trial)[46], registered with the Australian New Zealand Clinical Trials Registry (Registration number: ACTRN12618000732280). The use of plasma samples and subsequent data received ethical approval by the South Metropolitan Health Service Ethics Committee (Approval number: RGS731). Non-severe burns patients (n=20; total samples = 66) who suffered flame or scald burns that were less than 15% total burn surface area (TBSA), aged between 18 to 65 years of age and gave consent were recruited in the State Adult Burns Unit at Fiona Stanley Hospital (Western Australia) within 48 h of hospital admission. Collection of whole blood in lithium heparin tubes was scheduled on study enrolment (n=18) and the day of surgery (for burn wound debridement; n=10), as well as two days (n=11), two weeks (n=11), and six weeks post-surgery (n=16). Samples were centrifuged and stored at −80C within 4 hours of collection. At two weeks post-surgery, patient burn wounds were assessed for dressing cessation and stratified according to whether the wound was re-epithelialized or dressing continuation until wound closure (**Figure 1**). Mean (± SD) days from burn injury at the time of surgery was 6.1 (± 2.9) days, and at the six weeks post burns surgery timepoint was 45.64 (± 7.5) days, TBSA was 4.0% (± 3.6%) and age at time of collection was 39.87 (± 16.73) years. Additional demographic information of the non-severe burn cohort is tabulated in **Table 1**.

**Figure 1.**
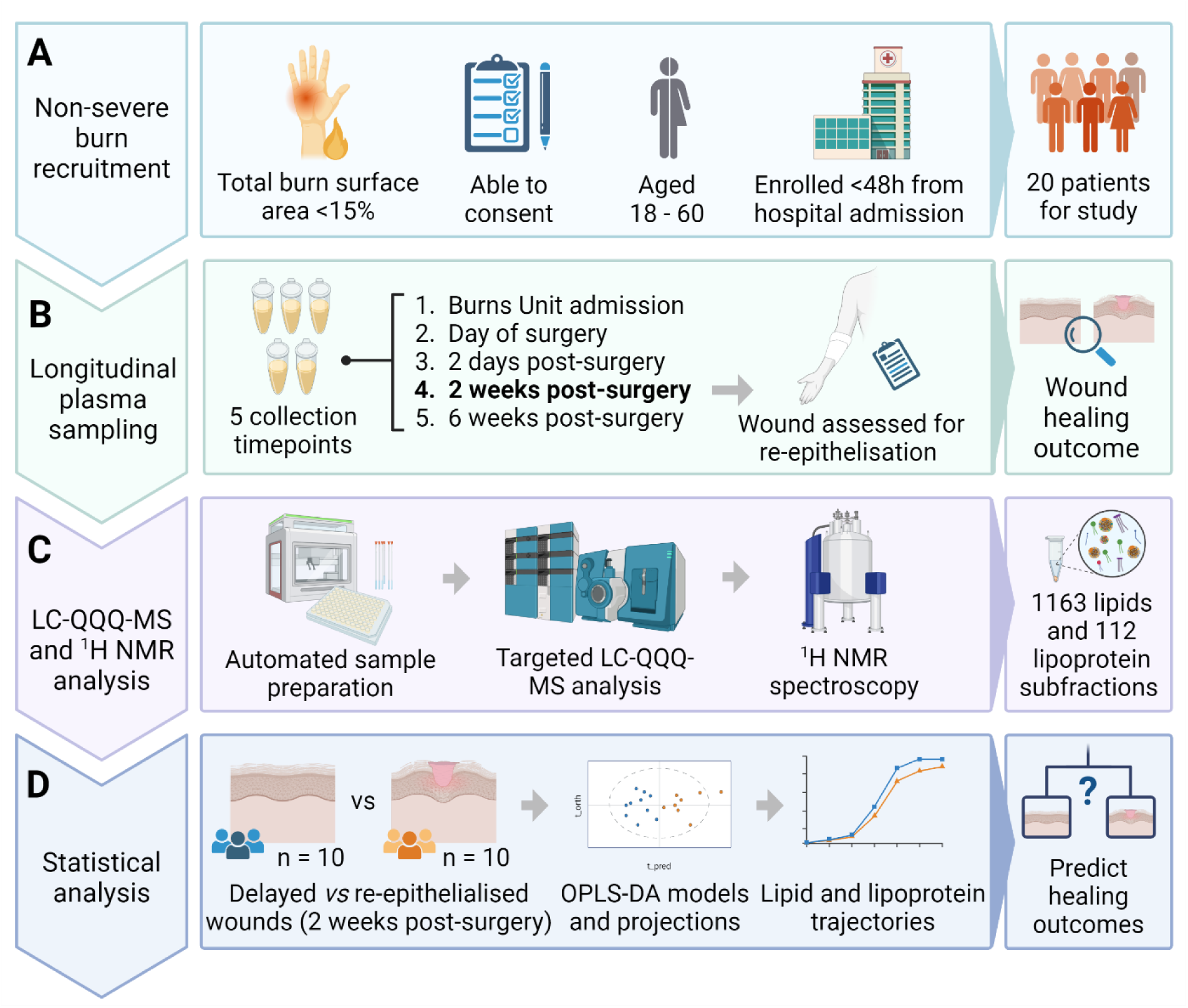
Generalised workflow of patient recruitment to the CABIN Fever trial, sample collection, lipid profiling, ^1^H NMR spectroscopy and statistical analysis for the study. **A)** Enrolment criteria for patients with non-severe burns into the CABIN Fever study, with final patient number used in this study (n=20). **B)** Longitudinal plasma sampling of patients, with plasma collected at trial enrolment, day of wound debridement surgery, two days, two weeks and six weeks post-surgery and burn wounds assessed at two weeks post for re-epithelisation. **C)** Simplified lipid profiling analysis using LC-QQQ-MS to detect 1163 lipid species spanning 20 subclasses and ^1^H NMR spectroscopy using 1D experiments to measure 112 lipoprotein subfractions. **D)** Statistical analysis comparing lipid profiles of patients with wound re-epithelisation (RE) (n=10) and delayed wound re-epithelisation (DRE) (n=10) at two weeks post-surgery using OPLS-DA modelling and lipid trajectories to predict wound healing outcomes pre-surgery. Image created using BioRender.com.

**Table 1.** Demographics and outcome parameters collected of the total non-severe burn patients recruited into the CABIN Fever trial, divided into overall and re-epithelisation groups. For each characteristic, p-values were computed using Wilcoxon rank sum test to observe any demographic differences.

| **Characteristic** | **Overall, N = 20<sup>1</sup>** | **Re-epithelisation, N = 10<sup>1</sup>** | **Delayed re-epithelisation, N = 10<sup>1</sup>** | **p-value<sup>2</sup>** |
| --- | --- | --- | --- | --- |
| **Sex** |  |  |  | >0.9 |
| *Male* | 17 (85%) | 9 (90%) | 8 (80%) |  |
| *Female* | 3 (15%) | 1(10%) | 2 (20%) |  |
| **Age** | 40 (25, 50) | 34 (26, 45) | 42 (25, 57) | 0.6 |
| **Days from burn injury to enrolment** | 2.70 (1.89, 3.98) | 2.91 (2.50, 4.27) | 2.03 (1.82, 3.72) | 0.4 |
| **TBSA %** | 3.55 (1.43, 6.13) | 1.85 (0.70, 4.20) | 5.05 (2.86, 6.54) | 0.089 |
| **Burn source** |  |  |  | 0.057 |
| *Flame* | 13 (65%) | 4 (40%) | 9 (90%) |  |
| *Scald* | 7 (35%) | 6 (60%) | 1 (10%) |  |
| **First aid before admission** | 14 (70%) | 8 (80%) | 6 (60%) | 0.6 |
| **Hospital length of stay in days** | 5 (2.75, 7) | 3 (1.25, 6.50) | 5 (5, 7) | 0.12 |
| **POSAS scores** | 35 (27, 42) | 30 (26, 42) | 38 (27, 40) | 0.5 |
| **Post-surgery in hospital days** | 3 (0, 4) | 0.50 (0, 2.50) | 3 (3, 4) | 0.092 |
| **Infection present** | 1 (5.0%) | 0 (0%) | 1 (10%) | >0.9 |
| **Antibiotics given** | 2 (10%) | 1 (10%) | 1 (10%) | >0.9 |
| **Surgery type** |  |  |  | >0.9 |
| *Graft* | 11 (55%) | 5 (50%) | 6 (60%) |  |
| *Cell autograft* | 6 (30%) | 3 (30%) | 3 (30%) |  |
| *Non-operative* | 3 (15%) | 2 (20%) | 1 (10%) |  |
| **Scars requiring laser** | 1 (5.0%) | 0 (0%) | 1 (10%) | >0.9 |
| **Re-graft required** | 1 (5.3%) | 0 (0%) | 1 (11%) | >0.9 |
| **Fitzpatrick skin phenotype** |  |  |  | 0.7 |
| *I* | 11 (55%) | 7 (70%) | 4 (40%) |  |
| *II* | 3 (15%) | 1 (10%) | 2 (20%) |  |
| *III* | 3 (15%) | 1 (10%) | 2 (20%) |  |
| *IV* | 2 (10%) | 1 (10%) | 1 (10%) |  |

| V | 1 (5.0%) | 0 (0%) | 1 (10%) |
| --- | --- | --- | --- |
| <sup>1</sup> n (%); Median (IQR) |  |  |  |
| <sup>2</sup> Fisher's exact test; Wilcoxon rank sum test; Wilcoxon rank sum exact test |  |  |  |

Abbreviations: TBSA – Total body surface area, POSAS – Patient and observer scar assessment scale.

### Pooled quality control samples

To ensure reproducibility and quality during the analytical runs, commercially available pooled human plasma (BioIVT, Westbury, NY, USA) was purchased and sub-aliquoted prior to LC-QQQ-MS and ^1^H NMR spectroscopy analysis. Pooled plasma samples were used as quality control (QC) samples throughout the runs between every tenth study sample.

### Lipid profiling

A comprehensive targeted lipid profiling approach was employed using a previously published LC-QQQ-MS method[47] to detect 1163 lipid species from 20 lipid subclasses in plasma and was used without any modifications (**Figure 1**). Briefly, using a Biomek i5 liquid handling system (Beckman Coulter, Mount Waverley, Victoria 3149, Australia) for automation, 90 µL of stable isotope labelled (SIL) internal standards diluted in isopropyl alcohol (IPA) was added to 10 µL of plasma. Following chilling and centrifuging at 14 000 *g* for 15 mins, 50 µL of supernatant was transferred to a 350 µL 96-well plate (Eppendorf, Macquarie Park, NSW) for same-day LC-QQQ-MS analysis. Targeted lipid profiling was performed on a ExionLC^TM^ coupled to a 6500+ QTRAP (SCIEX, Concord, CA).

### ^1^H NMR spectroscopy

^1^H NMR spectroscopy analysis of plasma samples was conducted using a published method[48]. Briefly, plasma samples were centrifuged, and the supernatant combined 1:1 with phosphate buffer (75 mM Na_2_HPO_4_, 2 mM NaN_3_, and 4.6 mM sodium trimethylsilyl propionate-[2,2,3,3-2H_4_] (TSP) in D_2_O, pH 7.4 ± 0.1) and mixed. 600 µL was then transferred into 5 mm outer diameter SampleJet NMR tubes and sealed with POM balls. Analysis was completed a 600 MHz Bruker Avance III HD spectrometer (Bruker BioSpin, Billerica, Massachusetts, USA) using the Bruker *in vitro* Diagnostics research (IVDr) methods. A standard ^1^H 1D experiment with solvent pre-saturation was run (32 scans, 98304 data points, 18028.85 Hz spectral width)[49] with an acquisition time of 4 min. Subsequently, a 4 min diffusion and relaxation editing (DIRE) experiment acquired 98 304 data points in 64 scans with a spectral width of 18 028.846 Hz[32,50,51].

### Lipoprotein subfraction analysis

The Bruker IVDr Lipoprotein Subclass Analysis (B.I.-LISA™) generates 112 lipoprotein parameters for each plasma sample using the standard ^1^H 1D experiment. This quantifies the −CH_2_ peak at δ = 1.25 and the −CH_3_ peak at δ = 0.80 in the normalised 1D spectrum (completed within Bruker QuantRef manager in Topspin) using a partial least squares (PLS-2) regression model [52]. The lipoprotein classes assessed included very low-density lipoproteins (VLDLs), low-density lipoproteins (LDLs), intermediate-density lipoproteins (IDLs), and high-density lipoproteins (HDLs). VLDL subfractions were categorised into six density classes (0.950-1.006 kg/L) as VLDL-1, VLDL-2, VLDL-3, VLDL-4, VLDL-5, and VLDL-6. LDL subfractions were assigned to six density classes as LDL-1 (1.019-1.031 kg/L), LDL-2 (1.031-1.034 kg/L), LDL-3 (1.034-1.037 kg/L), LDL-4 (1.037-1.040 kg/L), LDL-5 (1.040-1.044 kg/L), and LDL-6 (1.044-1.063 kg/L). HDL subfractions were classified into four density classes as HDL-1 (1.063-1.100 kg/L), HDL-2 (1.100-1.125 kg/L), HDL-3 (1.125-1.175 kg/L), and HDL-4 (1.175-1.210 kg/L). Additional details on lipoprotein analysis can be found in the Supplementary Information (**Table S1**).

Furthermore, the DIRE experiment was used to obtain GlycA (composite peak of *N*-acetyl signals from α-1-acid glycoprotein, α-1-antichymotrypsin, α-1-antitrypsin, haptoglobin, and transferrin) and GlycB (acetyl signal arising from glycoprotein *N*-acetylneuraminidino groups) signals. Pre-processing was comprised of baseline correction with an asymmetric least-squares routine and normalisation to the erectic signal using the *metabom8* version 1.0.0 R package (github.com/tkimhofer/metabom8). For this study, only GlycB was integrated (δ2.07) from the processed spectra as the GlycA signal (δ2.03) was overlapped by triacylglycerol signals, impeding results.

### Mass Spectrometry data integration

Raw spectra were imported into Skyline 21.1 (MacCoss, WA, USA) software for peak integration and data pre-processing and filtering was performed in R (version 4.4.1, r foundation, Vienna, Austria) in R Studio[53] (version 1.4.1, R Studio, Boston, MA, USA). Data pre-processing consisted of samples/lipid species with >50% missing values being filtered out, imputation of minimum value/2 in samples/lipid species with <50% missing values and filtering features with >30% relative standard deviations in plasma QC samples. Each feature underwent signal drift correction using the Random Forrest method in the *statTarget* R package[54] with pooled plasma QC samples.

### Statistical analysis

Principal component analysis (PCA), orthogonal projections to latent structures-discriminant analysis (OPLS-DA) models and projections were created using the *metabom8* R package (V0.2) from https://github.com/tkimhofer (**Figure 1**). Additionally using *metabom8*, the eruption plot[55] was produced by calculating univariate effect size, Cliff’s delta, on the x-axis and the OPLS-DA predictive loadings on the y-axis, with each point coloured by Mann-Whitney U test p-value. The training OPLS-DA model using samples from the time of burn admission was cross-validated using the Monte Carlo method on 2/3 of the dataset and 1/3 used to predict, with k subsets (for cross validation) set to 200. Univariate statistics and visualisation plots were generated in R (version 4.4.1, r foundation, Vienna, Austria) in R Studio[53] using the *ggplot2* (V3.4.2) R package from https://github.com/tidyverse/ggplot2.

## Results

### Data robustness

From pre-processing data acquired from LC-QQQ-MS, 850 lipid species passed the quality control checks of <50% missing values and <30% relative standard deviations across the interspersed pooled QC plasma. The lipid profiling and lipoprotein datasets were combined and underwent PCA modelling of study samples (n=66) and QC plasma samples (n=18) to evaluate robustness (**Figure S1**). QC plasma samples were tightly clustered in principal components 1 and 2 indicating minimal variability compared with study samples subfractions across the analytical run.

### Clinical parameters

No statistically significant differences were observed between the two recovery groups at admission in sex, age and burn TBSA% but the delayed re-epithelialisation (DRE) did have a higher mean TBSA% than the re-epithelisation (RE) group (**Table 1**). The median days from injury was not significantly different between re-epithelisation groups at burns admission (**Figure S2**), however the participants who presented to the Burns Unit more than 6 days since the initial injury were in the RE group. It should be noted that it was not feasible to collect plasma at every scheduled time point for each of the non-severe burn patients (n=20) and a total of 66 samples were analysed in this study. A comprehensive list of de-identified patient numbers at each collection timepoint is provided in **Table S2**.

For inflammatory monitoring, clinical measurements of CRP were not statistically significantly different at admission for both groups, but the median levels of CRP were elevated in the DRE group when compared to the RE group (**Figure S3**). Additionally, an acute phase glycoprotein signal arising from glycoprotein acetyl residues, GlycB, was obtained from the NMR spectra and is a marker of systemic inflammation[56,57]. GlycB levels at admission were not statistically significantly different between RE and DRE groups (**Figure S3**). Although, median GlycB levels were higher in the RE group compared to the DRE.

### Plasma lipidome and lipoprotein panel predict 2 week wound re-epithelisation outcomes for non-severe burns at hospital admission

Lipid and lipoprotein data from the admission timepoint were used to construct an OPLS-DA (**Figure 2A**) model for classification of RE *vs* DRE at the 2-week clinic visit. The model parameters included R^2^X (total variance) = 0.11, area under receiver operating characteristics (AUROC) = 1, and cross validation-AUROC (CV-AUROC) = 0.75 using the Monte Carlo method based on 2/3 of the dataset as the training set and the remainder as predictor, with k (number of model iterations) set to 200. For the projected models, the day of surgery had a sensitivity of 0.67 and a specificity of 1.0 (**Figure 2B**) using a confusion matrix for predicting wound recovery outcomes, 2 days post-surgery (**Figure 2C**; sensitivity = 0.83, specificity = 1.0), 2 weeks post-surgery (**Figure 2D**; sensitivity = 1.0, specificity = 0.67), and 6 weeks post-surgery (**Figure 2E**; sensitivity = 1.0, specificity = 0.75).

**Figure 2.**
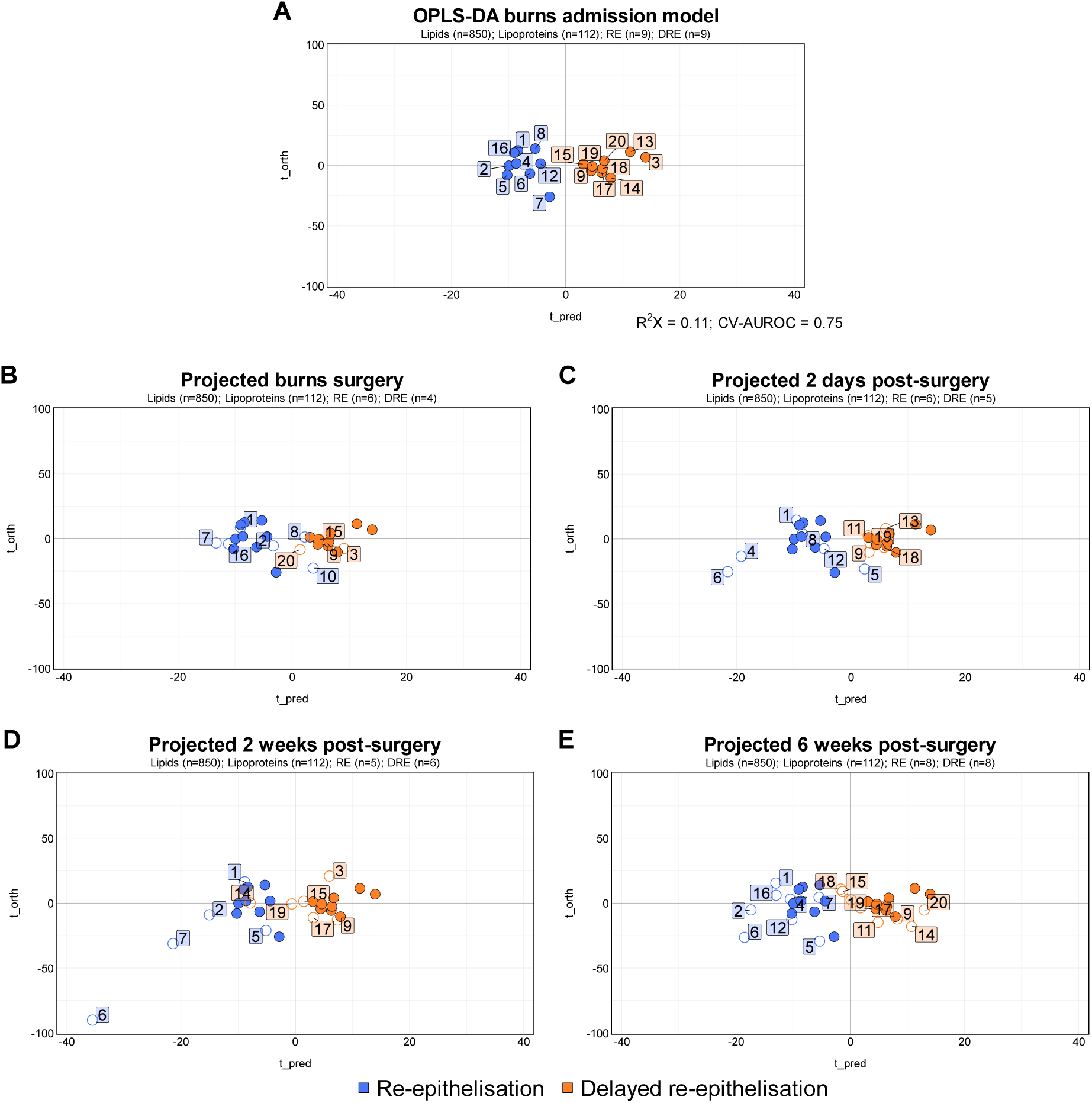
Supervised orthogonal projection to latent structures-discriminant analysis (OPLS-DA) modelling of re-epithelialisation and delayed re-epithelisation groups at burn admission, with projected timepoints of surgery, two days, two weeks and six weeks post-surgery. A) OPLS-DA model of admission (R^2^X = 0.11, AUROC = 1, CV-AUROC = 0.75) comparing RE (n=9; coloured blue) and DRE (n=9; coloured orange) using 850 lipid species and 112 lipoprotein subfractions. Each sequential plot (B-E) has predicted OPLS-DA scores of each timepoint projected on the admission model (coloured open circle) with an identifying patient number and sample sizes for RE and DRE groups are provided in the subtitle of each plot.

To visualise OPLS-DA loadings, an eruption plot[55] was produced with significant lipid and lipoproteins coloured by Mann-Whitney U p-value (red for p-values < 0.05; **Figure 3**). For patients with RE at two weeks post-surgery, key lipid species influencing the admission model were from the triacylglycerol (TG) and diacylglycerol (DG) subclasses, with phosphatidylglycerol (PG) PG(20:0_20:3) having the most influence (p-value = 0.012, Cliff’s delta = −0.70, OPLS-DA predictive component = 0.056), followed by DG(18:2_18:3) (p-value = 0.019, Cliff’s delta = −0.65, OPLS-DA predictive component = 0.051). Within the fatty acyl (FA) sidechains of the significant TGs and DGs, a recurring trend of FA(18:2), FA(18:3) and FA(20:4) was seen. In the DRE wound group, the main lipid subclasses driving separation were phosphatidylinositols (PIs), phosphatidylcholines (PCs) and phosphatidylserines (PSs), with PC(16:0_18:1) (p-value = 0.009, Cliff’s delta = 0.73, OPLS-DA predictive component = 0.81) and PC(16:0_16:0) (p-value=0.007, Cliff’s delta = 0.75, OPLS-DA predictive component = 0.078) showing the greatest influence on the model.

**Figure 3.**
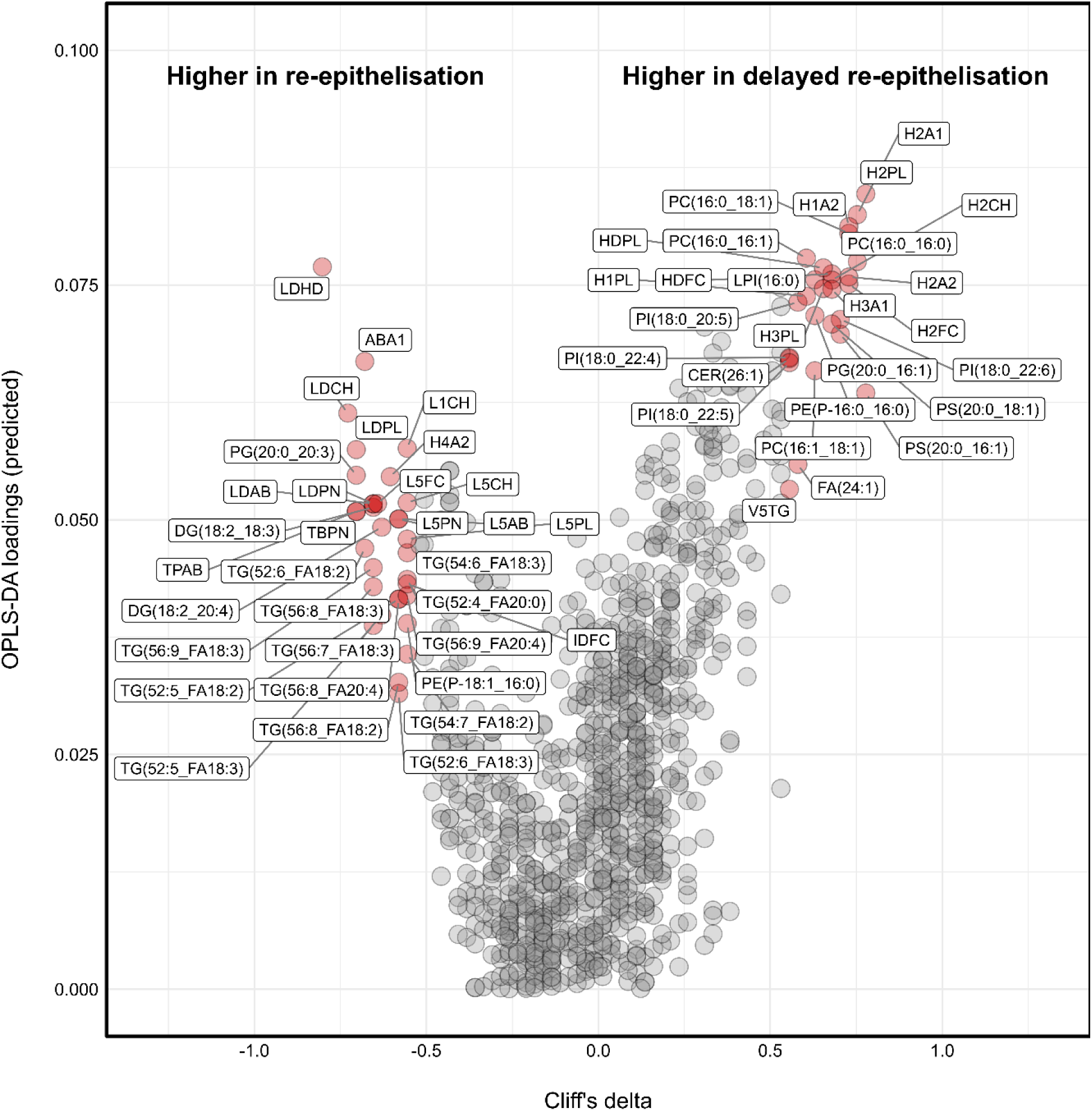
Eruption plot of the modelling of re-epithelisation versus delayed re-epithelisation groups at admission. Significant lipid species and lipoprotein subfractions from Mann-Whitney U analysis are highlighted in red (p-value <0.05) and corresponding Cliff’s delta direction for re-epithelisation groups are labelled (left – RE; right – DRE).

For lipoproteins, the RE group had increased levels of low-density lipoprotein (LDL) subfractions, with the ratio between LDL/HDL (LDHD) exerting the most influence (p-value = 0.004, Cliff’s delta = - 0.80, OPLS-DA predictive component = 0.077), followed by the apolipoprotein-B100/apolipoprotein-A1 (ABA1) ratio (p-value = 0.015, Cliff’s delta = −0.7, OPLS-DA predictive component = 0.067). In contrast, the DRE group showed higher levels of high-density lipoprotein (HDL) subfractions, particularly HDL subfraction 2 apolipoprotein-A1 (H2A1) (p-value = 0.007, Cliff’s delta = 0.75, OPLS-DA predictive component = 0.083) and HDL-2 phospholipids (H2PL) (p-value = 0.005, Cliff’s delta = 0.78, OPLS-DA predictive component = 0.085). On the contrary,

### Model validation using longitudinal data

To validate the model, data acquired from samples collected at surgery (**Figure 2B**), two days (**Figure 2C**), two weeks and six weeks post-surgery (**Figure 2E**) were projected onto the classification model. Resultant scores plots correctly predicted the DRE and RE patient groups regardless of timepoints, indicating a distinct multivariate signature of RE and DRE that is consistent over six weeks. The exception was three participants with DRE who moved closer to the RE space at the six weeks post-surgery timepoint and one at two weeks post, indicating a shift in their lipid and lipoprotein signature. Additionally, two RE participants clustered with the DRE group at surgery and one at two days post-surgery. Further investigation into the clinical outcome of the four incorrectly classified participants revealed no clinical abnormalities except for participant 14 who experienced an adverse inflammatory event independent to the burn.

### Differing lipidomic and lipoprotein signatures of re-epithelisation groups from burns admission to six weeks post-injury

To observe longitudinal trajectories of specific parameters, concentration values for the key lipids and lipoprotein subfractions highlighted in the eruption plot (Cliff’s delta > 0.5 or <-0.5 and p-value <0.05) (**Figure 3**) were plotted across the five timepoints (**Figure 4**). Concentrations of PC(16:0_16:0), PS(20:0_18:1), PI(18:0_20:5) and PS(20:0_16:1) were higher in the DRE patients compared to the RE group at nearly all timepoints except for surgery. Lysophosphatidylinositol (LPI) LPI(16:0) had similar patterns with the greatest difference in concentration at admission for DRE and RE, but converged at two days post-surgery with a subsequent increase in DRE above RE evolving over time. Lipoprotein subfractions H2A1, H2PL, HDL-1 apolipoprotein-A2 (H1A2), HDL-2 free cholesterol (H2FC) and HDL-2 apolipoprotein-A2 (H2A2) followed similar trends as the other lipid species, being higher in DRE at all timepoints except for H2FC that converges at surgery.

**Figure 4.**
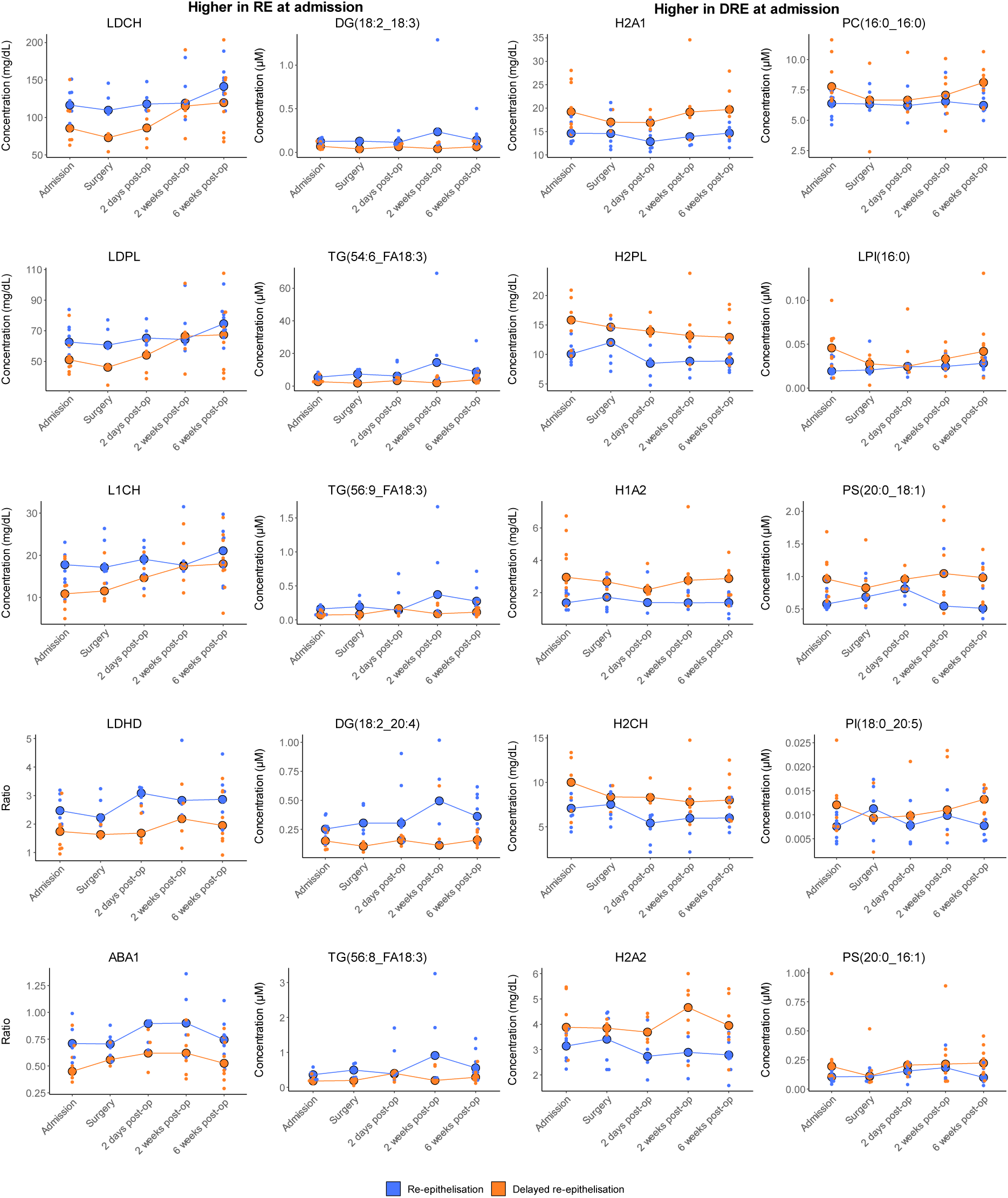
Trajectory dot plots of five significant and main separation driver lipids and lipoprotein subfractions from the eruption plot and OPLS-DA modelling for both wound outcome groups over time. Plots display lipid and lipoprotein subfraction individual concentrations of each patient as well as the median concentration with a trendline of both RE (n=10; coloured blue) and DRE (n=10; coloured orange) at burn admission, surgery, two days, two weeks, and six weeks post-surgery.

RE lipid trajectories for DG(18:2_18:3), TG(54:6_FA18:3), TG(56:9_FA18:3), TG(56:8_FA18:3) and DG(18:2_20:4) were similar, having higher concentrations than the RE but converging at two days post-surgery and spiking at two weeks post-surgery before returning to a similar concentration as at admission. For the lipoprotein subfractions, concentrations and ratios all were higher at all time points for LDHD, ABA1, LDL cholesterol (LDCH), LDL phospholipids (LDPL) and LDL subfraction 1 cholesterol (L1CH), except for LDPL and L1CH that had similar median concentrations in DRE at two weeks post-surgery. Interestingly, trajectories for both wound outcome groups for the significantly different lipid species had comparable median concentrations between their own admission and six weeks post-surgery.

Even though PC(16:0_16:1), PC(16:0_18:1), PG(20:0_16:1), PG(20:0_20:3), PI(18:0_22:6) and PC(16:1_18:1) lipid species were considered significant from univariate Mann-Whitney U test and had stronger effect sizes at admission (**Figure 3**), trajectories over time for both re-epithelisation groups appeared to be random (**Figure S4**) indicating these lipids may only be useful at burns admission for predicting wound healing and not post-surgery.

## Discussion

Using samples collected at hospital admission following a burn injury, we were able to prospectively classify individuals who experienced delayed re-epithelialisation (DRE) and required extra wound care from patients who had wound re-epithelialisation (RE) completed at two weeks post burn surgery.

RE and DRE wound outcome groups had distinct lipid and lipoprotein signatures at admission, as observed in the separation of the two groups in the OPLS-DA model (**Figure 2A**). At surgery and two days post-surgery (**Figure 2B, 2C**), the OPLS-DA model was able to predict both groups accurately, except for patient 5, 8, 10 in the RE group who clustered with the DRE. This could infer that the debridement surgery of their burn wounds may have altered the lipid and lipoproteins signatures in these patients to reflect more of the signatures seen in DRE in the short-term. However, these patients re-clustered with the RE group at two weeks post-surgery (**Figure 2C**, **Figure 4**). For all patients, with exception of participant 14, the predictive model accuracy for two weeks post-surgery was similar to the previous timepoints.

In contrast, patient 14 from the DRE group moving into the RE space which corresponded to an adverse inflammatory event experienced that was independent of the burn injury. This supports that the lipid and lipoprotein profiles observed in this study are influenced by inflammatory processes that are occurring. At six weeks post-surgery, the model was still able to accurately classify both groups, except for DRE patients 18 and 15 profiles moving towards the RE side. The accuracy of the OPLS-DA model and consistent separation of RE and DRE group infers that patients who experienced delayed wound closure at two weeks post-surgery had a distinct lipid and lipoprotein profile that occurred shortly after the initial burn injury and persisted up to six weeks post-surgery.

For the RE group, lipid and lipoprotein profiles may be indicating a divergent systemic response to injury and an underlying lipid-mediated inflammatory mechanism compared to the DRE group (**Figure 2**, **Figure 4**). GlycB is a composite signal from the 5-*N*-acetyl methyl groups of sialic acid in glycosylated acute phase protein markers during inflammation[58]. GlycB and CRP can correlate[58] but numerous studies have identified that the two markers can reflect different inflammatory processes[59], as seen in conditions such as obesity[60], type 2 diabetes[61] and cardiovascular disease[59]. With the difference observed in GlycB and CRP levels (though not statistically significant) between RE and DRE (**Figure S3**) two differing inflammatory patterns may have triggered in response to the burn injury. This is exemplified in the difference seen in the lipid and lipoprotein profiles between healing groups. Patients with wound closure at two weeks post-surgery had lipid species predominantly from the TG and DG subclasses with FA sidechains of 18:2, 18:3 and 20:4 and LDL lipoprotein subfractions (**Figure 3**). Analysing the trajectories of the most significant differentiating TGs and LDL subfractions (p-value < 0.02), the RE group had higher concentrations than the DRE group across all timepoints with a spike at two weeks post-surgery for some lipid species (**Figure 4**). The unique lipid and lipoprotein signature in the RE group may be reflecting cholesterol metabolism that promotes wound healing post-injury during inflammation. To our knowledge, no research has explored the role of TGs or DGs in the wound healing process. However, a recent study has established that in inflammatory conditions, circulating arachidonic acid (FA20:4), a precursor to the biosynthesis of inflammatory lipid mediators, can be re-esterified into TG species[62]. This re-esterification was associated with increasing levels of plasma LDL cholesterol (LDCH), apolipoprotein B100 and total cholesterol, inferring that arachidonic acid can regulate lipid and lipoprotein metabolism through influencing genes and proteins related to cholesterol metabolism[62]. At admission, patients within the RE group had higher levels of LDCH, ABA1, and TG and DG species with incorporated FA20:4, though inflammatory, may be highlighting on-going cholesterol metabolism that can assist wound healing processes. Cholesterol metabolism is important in the activation of the pro-inflammatory response as its influx into immune cells, such as B-cells, can trigger the activation of nuclear factor kappa β and mitogen-activated protein kinase signalling, leading to the expression of pro-inflammatory interleukin-6 and tumour necrosis factor-α cytokines [63,64]. Inflammation is integral to the wound healing process, facilitating necrotic tissue and cellular debris removal, pathogen clearance and stimulation of growth factors[65,66]. Although, persistent inflammation after this wound healing phase can be damaging and associated with poor patient outcomes particularly for severe burns[65]. As seen at 6 weeks post-injury (**Figure 2E**), this lipid and lipoprotein signature remains, which even though beneficial at admission for wound healing, could pre-dispose patients to long-term complications if it persists post-6 weeks.

Plasma lipid and lipoprotein profiles from non-severe burn patients with DRE were primarily comprised of phospholipids, specifically PSs, PIs and PCs (**Figure 3**). *In vivo*, phospholipids can inhibit pro-inflammatory processes to modulate inflammation[67], for example PSs after skin injury have been observed to bind to annexins on damaged cells to promote apoptotic cell clearance and coagulation during the healing process[68]. Additionally, higher concentrations of HDLs were observed, which are related to phospholipids, as the main constituent of HDL particles are phospholipids (60-70% by mass)[69]. The HDL phospholipid subfraction was also one of the main drivers of separation between the two groups in the model. The anti-inflammatory properties of HDLs, such as suppressing tumour necrosis factor-α, have been extensively researched in cardiovascular disease[70]. For the DRE group, the systemic response is possibly prioritising regulation through up-regulating anti-inflammatory molecules to combat the initial insult from the burn injury. Though a crucial process, it may not be beneficial for promoting wound closure in the short-term. Parallel findings from the Glue Grant that aimed to characterise the genomic response post-injury (including burn trauma) established that patients with co-morbidities post-injury had suppressed activation of protective immunity genes, particularly the interferon pathway, when compared to patients with no-complications post-injury[71]. Those with co-morbidities experienced worsened patient outcomes, such as longer times to recovery (>2 weeks post-injury) than the no-complications cohort[71], in which this dampened inflammatory gene response may be associated with. However, the findings for the DRE group may also suggest a higher metabolic demand due to the burn injury, resulting in lower levels of cholesterol-based lipoproteins and lipids compared to the RE group. The consumption of crucial cholesterol and lipid intermediates might predispose patients to poorer healing outcomes, such as delayed wound closure. Further investigation is needed to determine the observed metabolic effects in the DRE group and how these effects are associated with delayed wound closure at 2 weeks post-injury (**Figure 2D**).

The response in the DRE group could possibly be reflective of a phenomenon termed ‘burn wound conversion’. Burn wound conversion refers to the process by which a partial-thickness burn evolves into deeper dermal-thickness burns over time, leading to slower healing rates and necessitating surgical intervention[72]. The pathogenesis of burn wound conversion remains poorly understood; however, it is recognised that various factors contribute to it, including infection, improper inflammatory response, wound oedema, excessive eschar (dead tissue trapping healthy skin), and tissue desiccation (dryness)[72]. Wound conversion can commence as early as one hour post-injury and progress to full-thickness burns within 24 hours, depending on the severity of the initial burn[73]. Potentially, the DRE group may have undergone wound conversion prior to hospitalisation (>24 hours since injury; **Table 1**) and contributing to the distinctive lipid and lipoprotein signature observed at admission. However, data related to burn wound conversion were not available for this study and would require thorough validation and further research to support these findings.

In a study we previously conducted with the same non-severe burn cohort[33], we identified that non-severe burn injury may induce an inflammatory lipid and lipoprotein signature that persisted from admission to six weeks post-surgery when compared to non-burn controls. This initial study was crucial in providing insight into the impact on lipid and lipoprotein metabolism induced by the burn injury and underlying inflammatory mechanisms. Due to this, we found that the lipid species and lipoprotein subfractions that were main drivers of separation between non-severe burns and non-burn controls minimally overlapped with those observed between DRE and RE wound groups, confirming that these profiles are more associated with the wound healing process. With this information, clinical stratification and prediction for wound healing outcomes is more attainable as the OPLS-DA model that is based on burn admission shows a DRE lipid and lipoprotein signature that is not solely due to the impact of the initial injury.

Currently, monitoring burn wound healing is challenging as there is no standardised quantitative tool that can measure healing parameters. Assessment of burn wounds relies mostly on visual observation by clinicians, which can include calculating percentage of the total body surface area (TBSA) burn from charts and depth of injury that is mainly determined by the skin colour and elasticity[74]. As this is visual, there is a degree of subjectivity between clinicians even when specialised and trained, with a study reporting accuracy in burn wound assessment only 60-70% of the time[9]. Furthermore, wound dressings tend to be left on the skin between 2 to 5 days to facilitate healing, which limits opportunities for visual assessment[8]. New studies in non-severe burns have presented technologies that have potential to monitor burn wounds and predict patient outcomes regardless of dressing application[8,75]. Kenworthy et al. have reported that bioimpedance spectroscopy (BIS), the measure of the body’s inter-compartmental fluid volumes, correlated with healing rates in limb burns and is non-invasive[8]. Similarly, Khani et al. used wavelet Shannon entropy of terahertz time-domain waveforms to measure the rate of re-epithelisation with high accuracy in predicting wound healing outcomes in pigs[75]. However, these studies were conducted on the local wound rather than using biofluids, in which analysing biofluids, even though invasive, may be more reflective of wound healing outcomes as burn injuries cause a systemic response and numerous processes are involved in tissue repair[11,41]. Predictive modelling using metabolic phenotyping has had success in clinical translation, such as a recent study has developed an interactive framework that can accurately predict SARS-CoV-2 infection recoveries and stratify patients into risk groups based on their metabolic signatures[38], where we have demonstrated similar capability in early prediction of wound healing outcomes for non-severe burns.

Monitoring plasma lipid and lipoprotein profiles after a non-severe burn from admission to hospital could be valuable for predicting wound healing outcomes and guiding treatment options[76]. Patients who experienced DRE at two weeks post-surgery presented with differing plasma profiles from patients with RE, with profiles dominated by of phospholipids and HDL subfractions instead of TGs, DGs and LDL subfractions (**Figure 2, 3**). Patients at risk of poor re-epithelisation outcomes from hospital admission may benefit from delivery of bioactive molecules into the wound microenvironment, circumventing the divergent systemic responses occurring to locally promote wound closure[78]. Numerous studies have shown significant improvements in healing post-injury in animal models with dressings infused with different bioactive molecules, such as alginate from seaweed[79], silicate bioactive glass[80], thrombomodulin[81] and arginine derivatives[82]. With the time related information shown in lipid profiles in **Figure 4**, it is apparent that the phospholipids and HDL subfraction predominant lipid and lipoprotein profiles in the delayed re-epithelisation group remained constant after surgical wound debridement and became more pronounced at two weeks post-surgery.

This could infer that if these lipid profiles at admission remain constant after surgery that potentially other wound-based treatments could intervene earlier than two weeks post-surgery to improve healing outcomes. The proof of concept shows promise in using lipid and lipoprotein signatures to predict healing outcomes as early as admission and will require further validation in larger non-severe burn cohorts to gain power for clinical translation.

### Study limitations

Due to the nature of the non-severe burn trial, we acknowledge there are limitations in this study mainly in sampling duration, sample size and distribution of sex. However, despite the relatively low sample size of the RE and DRE groups, particularly at surgery (**Figure 2, Table S1**), the lipid and lipoprotein panel that could differentiate RE from DRE remained constant across the timepoints, adding confidence that the observed differences in the lipidome was biologically relevant. These small sample sizes could result in type II errors and further research with larger sample sizes is needed to confirm the clinical differences observed in the lipid and lipoprotein profiles of the RE and DRE groups[83]. Furthermore, the trajectories cease at the 6-week post-surgery timepoint where the key influencing lipid and lipoprotein concentrations appeared to be returning to similar concentrations as seen at admission (**Figure 4**). Extending the timepoints past this 6-week mark may be beneficial when monitoring trajectories to visualise whether these lipid and lipoprotein profiles change after re-epithelisation of the DRE group and if they differ from hospital admission.

Lastly, the patients recruited into the trial were unequal in terms of sex distribution. To observe any effect of sex on the key influencing lipids and lipoproteins identified driving re-epithelisation groups (**Figure 4**), univariate tests (Mann-Whitney U) were applied to both sexes using lipid and lipoprotein concentrations at admission (the basis for the OPLS-DA model) (**Figure S5**). No significant differences were seen in the key lipid and lipoproteins, except for the ABA1 ratio where females had higher ABA1 ratios on average at admission (p-value = 0.035).

### Conclusion

At study enrolment, the plasma lipidome appeared to be prognostically different between patients who had re-epithelialised or who had delayed re-epithelisation of non-severe burn wounds at two weeks post-surgery. These distinctive profiles were maintained over the 6-week recovery period of monitoring with OPLS-DA modelling accurately predicting re-epithelisation outcomes at each of the sample collection timepoints. Additionally, the profiles of the inflammatory markers GlycB and CRP were distinct different at admission between the DRE and RE groups, suggesting divergent systemic responses to the burn injury. The differential lipid profiles of patients with DRE wounds had phospholipids, specifically PGs, PIs and PCs determined to be the key univariately significant lipid species when compared to the RE group. Of the lipoprotein subfractions, HDLs, particularly HDL-2 subfractions were drivers of separation for the DRE group. Opposing this, patients with RE wounds displayed lipid profiles dominated by mainly TGs and some DGs with univariately significant lipid species (TG(54:6_FA18:3), TG(56:9_FA18:3), TG(56:8_FA18:3), DG(18:2_18:3) and DG(18:2_20:4)). Additionally, LDL subfractions, such as LDCH and LDPL, were predominant in the lipoprotein profiles. This profile suggests a potential role of cholesterol metabolism in wound healing processes, but persistent inflammation beyond 6 weeks may lead to long-term complications. Furthermore, lipid and lipoprotein subfraction trajectories of the phospholipid species and HDL subfractions remained higher in the DRE than the RE group across all timepoints, potentially signifying the systemic response prioritising modulation of the inflammation induced post-burn that may not be beneficial for wound closure in the short-term. These findings exemplify the value in monitoring plasma lipid and lipoprotein signatures after a non-severe burn and that wound healing outcomes can potentially be predicted at admission to hospital but necessitates subsequent validation in larger cohorts of non-severe burns to establish clinical utility.

## Declarations

### Ethics approval

The randomised placebo-controlled trial of Celecoxib for Acute Burn Inflammation and fever was registered with the Australian New Zealand Clinical Trials Registry (Registration number: ACTRN12618000732280). Ethical approval for the use of the retrospectively collected samples for this study was granted by the South Metropolitan Health Service Health Research Ethics Committee (Approval number: RGS731).

### Availability of data and materials

The datasets used and/or analysed in the current study are available from the corresponding author upon request.

### Authors contributions

ER, MWF and FMW designed, led, and provided the samples for the CABIN Fever study. MR, GLM, ER, MWF, FMW, EH, JKN, LW and NG contributed to study concept and design. MR analysed the samples by mass spectrometry and drafted the manuscript. SL and PN analysed the samples by nuclear magnetic resonance spectroscopy. RM, SL, PN, JW and JKN curated the lipoprotein dataset and provided statistical/interpretation input. MR and LW performed statistical analysis of the combined dataset. MR, ER, GLM, MWF, FMW, RM, SL, PN, JW, EH, JKN, NG and LW edited and critically revised the manuscript. ER, FMW, EH and JKN provided funding support. All authors read and approved the final manuscript.

## Abbreviations

AUROC: Area under receiver operating characteristics
CABIN: Randomised placebo-controlled trial of Celecoxib for Acute Burn Inflammation and Fever
CRP: C-reactive protein
CV: Cross-validated
DIRE: Diffusion and relaxation editing
DG: Diacylglycerol
DRE: Delayed wound re-epithelisation
FA: Fatty acyl
HDL: High-density lipoprotein
IDL: Intermediate-density lipoprotein
IPA: Isopropyl alcohol
IQR: Inter-quartile range
LC-QQQ-MS: Liquid chromatography-tandem mass spectrometry
LDCH: Low-density lipoprotein cholesterol
LDHD: Low-density lipoprotein / high-density lipoprotein ratio
LDL: Low-density lipoprotein
LDPL: Low-density lipoprotein phospholipids
LPI: Lysophosphatidylinositol
NMR: Nuclear magnetic resonance
OPLS-DA: Orthogonal projections to latent structures-discriminant analysis
PC: Phosphotidylcholine
PCA: Principal component analysis
PG: Phosphatidylglycerol
PI: Phosphatidylinositol
PLS: Partial least squares
POSAS: Patient and observer scar assessment scale
PS: Phosphotidylserine
QC: Quality control
QTRAP: Quadrupole ion trap
RE: Wound re-epithelisation
SD: Standard deviation
TBSA: Total body surface area
TG: Triacylglyerol
VLDL: Very low-density lipoprotein

## References

[1] Greenhalgh DG. Management of Burns. N Engl J Med 2019;380:2349–59. 10.1056/NEJMra1807442.

[2] Markiewicz-Gospodarek A, Kozioł M, Tobiasz M, Baj J, Radzikowska-Büchner E, Przekora A. Burn Wound Healing: Clinical Complications, Medical Care, Treatment, and Dressing Types: The Current State of Knowledge for Clinical Practice. International Journal of Environmental Research and Public Health 2022;19:1338. 10.3390/ijerph19031338.

[3] Stander M, Wallis LA. The Emergency Management and Treatment of Severe Burns. Emerg Med Int 2011;2011:161375. 10.1155/2011/161375.

[4] Wang Y, Beekman J, Hew J, Jackson S, Issler-Fisher AC, Parungao R, et al. Burn injury: Challenges and advances in burn wound healing, infection, pain and scarring. Advanced Drug Delivery Reviews 2018;123:3–17. 10.1016/j.addr.2017.09.018.

[5] Li Z, Maitz P. Cell therapy for severe burn wound healing. Burns Trauma 2018;6:13. 10.1186/s41038-018-0117-0.

[6] Osborne T, Edgar D, Gittings P, Wood F, Le Huray T, Allan B, et al. A prospective pilot study of the energy balance profiles in acute non-severe burn patients. Burns 2022;48:184–90. 10.1016/j.burns.2021.03.002.

[7] Shpichka A, Butnaru D, Bezrukov EA, Sukhanov RB, Atala A, Burdukovskii V, et al. Skin tissue regeneration for burn injury. Stem Cell Res Ther 2019;10:94. 10.1186/s13287-019-1203-3.

[8] Kenworthy P, Phillips M, Grisbrook TL, Gibson W, Wood FM, Edgar DW. Monitoring wound healing in minor burns—A novel approach. Burns 2018;44:70–6. 10.1016/j.burns.2017.06.007.

[9] Monstrey S, Hoeksema H, Verbelen J, Pirayesh A, Blondeel P. Assessment of burn depth and burn wound healing potential. Burns 2008;34:761–9. 10.1016/j.burns.2008.01.009.

[10] Li Z, Maitz P. Cell therapy for severe burn wound healing. Burns & Trauma 2018;6. 10.1186/s41038-018-0117-0.

[11] Wilkinson HN, Hardman MJ. Wound healing: cellular mechanisms and pathological outcomes. Open Biology 2020;10:200223. 10.1098/rsob.200223.

[12] Dhall S, Wijesinghe DS, Karim ZA, Castro A, Vemana HP, Khasawneh FT, et al. Arachidonic acid-derived signaling lipids and functions in impaired healing. Wound Repair Regen 2015;23:644–56. 10.1111/wrr.12337.

[13] Schierle CF, De la Garza M, Mustoe TA, Galiano RD. Staphylococcal biofilms impair wound healing by delaying reepithelialization in a murine cutaneous wound model. Wound Repair and Regeneration 2009;17:354–9. 10.1111/j.1524-475X.2009.00489.x.

[14] Plichta JK, Holmes CJ, Gamelli RL, Radek KA. Local Burn Injury Promotes Defects in the Epidermal Lipid and Antimicrobial Peptide Barriers in Human Autograft Skin and Burn Margin: Implications for Burn Wound Healing and Graft Survival. J Burn Care Res 2017;38:e212–26. 10.1097/BCR.0000000000000357.

[15] Duke JM, Randall SM, Wood FM, Boyd JH, Fear MW. Burns and long-term infectious disease morbidity: A population-based study. Burns 2017;43:273–81. 10.1016/j.burns.2016.10.020.

[16] Finnerty CC, Jeschke MG, Branski LK, Barret JP, Dziewulski P, Herndon DN. Hypertrophic scarring: the greatest unmet challenge after burn injury. Lancet 2016;388:1427–36. 10.1016/S0140-6736(16)31406-4.

[17] Frescos N. Assessment of pain in chronic wounds: A survey of Australian health care practitioners. International Wound Journal 2018;15:943–9. 10.1111/iwj.12951.

[18] Hamed K, Giles N, Anderson J, Phillips JK, Dawson LF, Drummond P, et al. Changes in cutaneous innervation in patients with chronic pain after burns. Burns 2011;37:631–7. 10.1016/j.burns.2010.11.010.

[19] Yoon J, Kym D, Hur J, Won JH, Yim H, Cho YS, et al. Time-varying discrimination accuracy of longitudinal biomarkers for the prediction of mortality compared to assessment at fixed time point in severe burns patients. BMC Emergency Medicine 2021;21:1. 10.1186/s12873-020-00394-z.

[20] Song J, Ozhathil DK, El Ayadi A, Golovko G, Wolf SE. C-reactive protein elevation is associated with increased morbidity and mortality in elderly burned patients. Burns 2023;49:806–12. 10.1016/j.burns.2022.05.008.

[21] Finnerty CC, Ju H, Spratt H, Victor S, Jeschke MG, Hegde S, et al. Proteomics Improves the Prediction of Burns Mortality: Results from Regression Spline Modeling. Clinical and Translational Science 2012;5:243–9. 10.1111/j.1752-8062.2012.00412.x.

[22] Muñoz B, Suárez-Sánchez R, Hernández-Hernández O, Franco-Cendejas R, Cortés H, Magaña JJ. From traditional biochemical signals to molecular markers for detection of sepsis after burn injuries. Burns 2019;45:16–31. 10.1016/j.burns.2018.04.016.

[23] Rittenhouse BA, Rizzo JA, Shields BA, Rowan MP, Aden JK, Salinas J, et al. Predicting wound healing rates and survival with the use of automated serial evaluations of burn wounds. Burns 2019;45:48–53. 10.1016/j.burns.2018.10.018.

[24] Rowan MP, Cancio LC, Elster EA, Burmeister DM, Rose LF, Natesan S, et al. Burn wound healing and treatment: review and advancements. Crit Care 2015;19:243. 10.1186/s13054-015-0961-2.

[25] Fear VS, Poh W-P, Valvis S, Waithman JC, Foley B, Wood FM, et al. Timing of excision after a non-severe burn has a significant impact on the subsequent immune response in a murine model. Burns 2016;42:815–24. 10.1016/j.burns.2016.01.013.

[26] Xiao-Wu W, Herndon DN, Spies M, Sanford AP, Wolf SE. Effects of Delayed Wound Excision and Grafting in Severely Burned Children. Archives of Surgery 2002;137:1049–54. 10.1001/archsurg.137.9.1049.

[27] Mathur S, Sutton J. Personalized medicine could transform healthcare. Biomed Rep 2017;7:3–5. 10.3892/br.2017.922.

[28] Duke J, Wood F, Semmens J, Spilsbury K, Edgar DW, Hendrie D, et al. A 26-year population-based study of burn injury hospital admissions in Western Australia. J Burn Care Res 2011;32:379–86. 10.1097/BCR.0b013e318219d16c.

[29] Buergel T, Steinfeldt J, Ruyoga G, Pietzner M, Bizzarri D, Vojinovic D, et al. Metabolomic profiles predict individual multidisease outcomes. Nat Med 2022;28:2309–20. 10.1038/s41591-022-01980-3.

[30] Yau A, Fear MW, Gray N, Ryan M, Holmes E, Nicholson JK, et al. Enhancing the accuracy of surgical wound excision following burns trauma via application of Rapid Evaporative IonisationMass Spectrometry (REIMS). Burns 2022;48:1574–83. 10.1016/j.burns.2022.08.021.

[31] Gray N, Lawler NG, Zeng AX, Ryan M, Bong SH, Boughton BA, et al. Diagnostic potential of the plasma lipidome in infectious disease: Application to acute sars-cov-2 infection. Metabolites 2021;11. 10.3390/METABO11070467.

[32] Lodge S, Nitschke P, Kimhofer T, Wist J, Bong S-H, Loo RL, et al. Diffusion and Relaxation Edited Proton NMR Spectroscopy of Plasma Reveals a High-Fidelity Supramolecular Biomarker Signature of SARS-CoV-2 Infection. Anal Chem 2021;93:3976–86. 10.1021/acs.analchem.0c04952.

[33] Ryan MJ, Raby E, Whiley L, Masuda R, Lodge S, Nitschke P, et al. Nonsevere Burn Induces a Prolonged Systemic Metabolic Phenotype Indicative of a Persistent Inflammatory Response Postinjury. J Proteome Res 2023. 10.1021/acs.jproteome.3c00516.

[34] Tzoulaki I, Castagné R, Boulangé CL, Karaman I, Chekmeneva E, Evangelou E, et al. Serum metabolic signatures of coronary and carotid atherosclerosis and subsequent cardiovascular disease. European Heart Journal 2019;40:2883–96. 10.1093/eurheartj/ehz235.

[35] Fromentin S, Forslund SK, Chechi K, Aron-Wisnewsky J, Chakaroun R, Nielsen T, et al. Microbiome and metabolome features of the cardiometabolic disease spectrum. Nat Med 2022;28:303–14. 10.1038/s41591-022-01688-4.

[36] Sun Y, Zou H, Li X, Xu S, Liu C. Plasma Metabolomics Reveals Metabolic Profiling For Diabetic Retinopathy and Disease Progression. Frontiers in Endocrinology 2021;12.

[37] Masuda R, Lodge S, Nitschke P, Spraul M, Schaefer H, Bong S-H, et al. Integrative Modeling of Plasma Metabolic and Lipoprotein Biomarkers of SARS-CoV-2 Infection in Spanish and Australian COVID-19 Patient Cohorts. J Proteome Res 2021;20:4139–52. 10.1021/acs.jproteome.1c00458.

[38] Ruffieux H, Hanson AL, Lodge S, Lawler NG, Whiley L, Gray N, et al. A patient-centric modeling framework captures recovery from SARS-CoV-2 infection. Nat Immunol 2023;24:349–58. 10.1038/s41590-022-01380-2.

[39] Vande Voorde J, Steven RT, Najumudeen AK, Ford CA, Dexter A, Gonzalez-Fernandez A, et al. Metabolic profiling stratifies colorectal cancer and reveals adenosylhomocysteinase as a therapeutic target. Nat Metab 2023;5:1303–18. 10.1038/s42255-023-00857-0.

[40] Qi S, Wu Q, Chen Z, Zhang W, Zhou Y, Mao K, et al. High-resolution metabolomic biomarkers for lung cancer diagnosis and prognosis. Sci Rep 2021;11:11805. 10.1038/s41598-021-91276-2.

[41] Begum S, Johnson BZ, Morillon A-C, Yang R, Bong SH, Whiley L, et al. Systemic long-term metabolic effects of acute non-severe paediatric burn injury. Sci Rep 2022;12:13043. 10.1038/s41598-022-16886-w.

[42] Qi P, Abdullahi A, Stanojcic M, Patsouris D, Jeschke MG. Lipidomic analysis enables prediction of clinical outcomes in burn patients. Sci Rep 2016;6:38707. 10.1038/srep38707.

[43] Kamolz L-P, Andel H, Mittlböck M, Winter W, Haslik W, Meissl G, et al. Serum cholesterol and triglycerides: potential role in mortality prediction. Burns 2003;29:810–5. 10.1016/S0305-4179(03)00196-7.

[44] Bjerrum JT, Wang Y, Zhang J, Riis LB, Nielsen OH, Seidelin JB. Lipidomic Trajectories Characterize Delayed Mucosal Wound Healing in Quiescent Ulcerative Colitis and Identify Potential Novel Therapeutic Targets. International Journal of Biological Sciences 2022;18:1813–28. 10.7150/ijbs.67112.

[45] Ryan M, Raby E, Masuda R, Lodge S, Nitschke P, Maker G, et al. Metabolic consequences of non-severe burn injury are associated with increased plasma markers of inflammation and cardiovascular disease risk. 2023. 10.1101/2023.04.24.537960.

[46] Raby E, Gittings P, Litton E, Berghuber A, Edgar DW, Camilleri J, et al. Celecoxib to improve scar quality following acute burn injury: Lessons learned after premature termination of a randomised trial. Burns Open 2024;8:128–35. 10.1016/j.burnso.2024.03.001.

[47] Ryan MJ, Grant-St James A, Lawler NG, Fear MW, Raby E, Wood FM, et al. Comprehensive Lipidomic Workflow for Multicohort Population Phenotyping Using Stable Isotope Dilution Targeted Liquid Chromatography-Mass Spectrometry. J Proteome Res 2023. 10.1021/acs.jproteome.2c00682.

[48] Dona AC, Jiménez B, Schäfer H, Humpfer E, Spraul M, Lewis MR, et al. Precision High-Throughput Proton NMR Spectroscopy of Human Urine, Serum, and Plasma for Large-Scale Metabolic Phenotyping. Anal Chem 2014;86:9887–94. 10.1021/ac5025039.

[49] Nicholson JK, Foxall PJ, Spraul M, Farrant RD, Lindon JC. 750 MHz 1H and 1H-13C NMR spectroscopy of human blood plasma. Anal Chem 1995;67:793–811. 10.1021/ac00101a004.

[50] Liu M, Nicholson JK, Lindon JC. High-resolution diffusion and relaxation edited one- and two-dimensional 1H NMR spectroscopy of biological fluids. Anal Chem 1996;68:3370–6. 10.1021/ac960426p.

[51] Liu M, Nicholson JK, Parkinson JA, Lindon JC. Measurement of Biomolecular Diffusion Coefficients in Blood Plasma Using Two-Dimensional 1H−1H Diffusion-Edited Total-Correlation NMR Spectroscopy. Anal Chem 1997;69:1504–9. 10.1021/ac9612133.

[52] Jiménez B, Holmes E, Heude C, Tolson RF, Harvey N, Lodge SL, et al. Quantitative Lipoprotein Subclass and Low Molecular Weight Metabolite Analysis in Human Serum and Plasma by 1H NMR Spectroscopy in a Multilaboratory Trial. Anal Chem 2018;90:11962–71. 10.1021/acs.analchem.8b02412.

[53] R Core Team. R: A Language and Environment for Statistical Computing. 2022.

[54] Luan H, Ji F, Chen Y, Cai Z. statTarget: A streamlined tool for signal drift correction and interpretations of quantitative mass spectrometry-based omics data. Anal Chim Acta 2018;1036:66–72. 10.1016/j.aca.2018.08.002.

[55] Kimhofer T, Lodge S, Whiley L, Gray N, Loo RL, Lawler NG, et al. Integrative Modeling of Quantitative Plasma Lipoprotein, Metabolic, and Amino Acid Data Reveals a Multiorgan Pathological Signature of SARS-CoV-2 Infection. Journal of Proteome Research 2020;19:4442–54. 10.1021/ACS.JPROTEOME.0C00519.

[56] Nitschke P, Lodge S, Kimhofer T, Masuda R, Bong S-H, Hall D, et al. J-Edited DIffusional Proton Nuclear Magnetic Resonance Spectroscopic Measurement of Glycoprotein and Supramolecular Phospholipid Biomarkers of Inflammation in Human Serum. Anal Chem 2022;94:1333–41. 10.1021/acs.analchem.1c04576.

[57] Duprez DA, Jacobs Jr DR. GlycA, a composite low-grade inflammatory marker, predicts mortality: prime time for utilization? Journal of Internal Medicine 2019;286:610–2. 10.1111/joim.12961.

[58] Fung E, Chan EYS, Ng KH, Yu KM, Li H, Wang Y. Towards clinical application of GlycA and GlycB for early detection of inflammation associated with (pre)diabetes and cardiovascular disease: recent evidence and updates. J Inflamm (Lond) 2023;20:32. 10.1186/s12950-023-00358-7.

[59] Connelly MA, Otvos JD, Shalaurova I, Playford MP, Mehta NN. GlycA, a novel biomarker of systemic inflammation and cardiovascular disease risk. J Transl Med 2017;15:219. 10.1186/s12967-017-1321-6.

[60] Levine JA, Han JM, Wolska A, Wilson SR, Patel TP, Remaley AT, et al. Associations of GlycA and High-Sensitivity C-Reactive Protein with Measures of Lipolysis in Adults with Obesity. J Clin Lipidol 2020;14:667–74. 10.1016/j.jacl.2020.07.012.

[61] Fizelova M, Jauhiainen R, Kangas AJ, Soininen P, Ala-Korpela M, Kuusisto J, et al. Differential Associations of Inflammatory Markers With Insulin Sensitivity and Secretion: The Prospective METSIM Study. J Clin Endocrinol Metab 2017;102:3600–9. 10.1210/jc.2017-01057.

[62] Li F, Wang Y, Yu H, Gao X, Li L, Sun H, et al. Arachidonic acid is associated with dyslipidemia and cholesterol-related lipoprotein metabolism signatures. Front Cardiovasc Med 2022;9. 10.3389/fcvm.2022.1075421.

[63] Bauer R, Brüne B, Schmid T. Cholesterol metabolism in the regulation of inflammatory responses. Front Pharmacol 2023;14. 10.3389/fphar.2023.1121819.

[64] Li Y, Schwabe RF, DeVries-Seimon T, Yao PM, Gerbod-Giannone M-C, Tall AR, et al. Free cholesterol-loaded macrophages are an abundant source of tumor necrosis factor-alpha and interleukin-6: model of NF-kappaB- and map kinase-dependent inflammation in advanced atherosclerosis. J Biol Chem 2005;280:21763–72. 10.1074/jbc.M501759200.

[65] Korkmaz HI, Flokstra G, Waasdorp M, Pijpe A, Papendorp SG, de Jong E, et al. The Complexity of the Post-Burn Immune Response: An Overview of the Associated Local and Systemic Complications. Cells 2023;12:345. 10.3390/cells12030345.

[66] Barrientos S, Stojadinovic O, Golinko MS, Brem H, Tomic-Canic M. Growth factors and cytokines in wound healing. Wound Repair Regen 2008;16:585–601. 10.1111/j.1524-475X.2008.00410.x.

[67] Bretscher P, Egger J, Shamshiev A, Trötzmüller M, Köfeler H, Carreira EM, et al. Phospholipid oxidation generates potent anti-inflammatory lipid mediators that mimic structurally related pro-resolving eicosanoids by activating Nrf2. EMBO Molecular Medicine 2015;7:593–607. 10.15252/emmm.201404702.

[68] Kreft S, Klatt AR, Straßburger J, Pöschl E, Flower RJ, Eming S, et al. Skin Wound Repair Is Not Altered in the Absence of Endogenous AnxA1 or AnxA5, but Pharmacological Concentrations of AnxA4 and AnxA5 Inhibit Wound Hemostasis. Cells Tissues Organs 2016;201:287–98. 10.1159/000445106.

[69] Kontush A, Lhomme M, Chapman MJ. Unraveling the complexities of the HDL lipidome1. J Lipid Res 2013;54:2950–63. 10.1194/jlr.R036095.

[70] Jia C, Anderson JLC, Gruppen EG, Lei Y, Bakker SJL, Dullaart RPF, et al. High-Density Lipoprotein Anti-Inflammatory Capacity and Incident Cardiovascular Events. Circulation 2021;143:1935–45. 10.1161/CIRCULATIONAHA.120.050808.

[71] Tompkins RG. Genomics of Injury: The Glue Grant Experience. J Trauma Acute Care Surg 2015;78:671–86. 10.1097/TA.0000000000000568.

[72] Singh V, Devgan L, Bhat S, Milner SM. The pathogenesis of burn wound conversion. Ann Plast Surg 2007;59:109–15. 10.1097/01.sap.0000252065.90759.e6.

[73] Hirth D, McClain SA, Singer AJ, Clark RAF. Endothelial necrosis at 1 hour postburn predicts progression of tissue injury. Wound Repair and Regeneration 2013;21:563–70. 10.1111/wrr.12053.

[74] Flanagan M. Wound measurement: can it help us to monitor progression to healing? J Wound Care 2003;12:189–94. 10.12968/jowc.2003.12.5.26493.

[75] Khani ME, Osman OB, Harris ZB, Chen A, Zhou J-W, Singer AJ, et al. Accurate and early prediction of the wound healing outcome of burn injuries using the wavelet Shannon entropy of terahertz time-domain waveforms. JBO 2022;27:116001. 10.1117/1.JBO.27.11.116001.

[76] Meikle TG, Huynh K, Giles C, Meikle PJ. Clinical lipidomics: realizing the potential of lipid profiling. Journal of Lipid Research 2021;62:100127. 10.1016/j.jlr.2021.100127.

[77] Farina JA, Rosique MJ, Rosique RG. Curbing Inflammation in Burn Patients. Int J Inflam 2013;2013:715645. 10.1155/2013/715645.

[78] Li R, Liu K, Huang X, Li D, Ding J, Liu B, et al. Bioactive Materials Promote Wound Healing through Modulation of Cell Behaviors. Adv Sci (Weinh) 2022;9:e2105152. 10.1002/advs.202105152.

[79] Stenlund P, Enstedt L, Gilljam KM, Standoft S, Ahlinder A, Lundin Johnson M, et al. Development of an All-Marine 3D Printed Bioactive Hydrogel Dressing for Treatment of Hard-to-Heal Wounds. Polymers 2023;15:2627. 10.3390/polym15122627.

[80] Hu S, Chang J, Liu M, Ning C. Study on antibacterial effect of 45S5 Bioglass®. J Mater Sci: Mater Med 2009;20:281–6. 10.1007/s10856-008-3564-5.

[81] Hsu Y-Y, Liu K-L, Yeh H-H, Lin H-R, Wu H-L, Tsai J-C. Sustained release of recombinant thrombomodulin from cross-linked gelatin/hyaluronic acid hydrogels potentiate wound healing in diabetic mice. European Journal of Pharmaceutics and Biopharmaceutics 2019;135:61–71. 10.1016/j.ejpb.2018.12.007.

[82] Zhang S, Hou J, Yuan Q, Xin P, Cheng H, Gu Z, et al. Arginine derivatives assist dopamine-hyaluronic acid hybrid hydrogels to have enhanced antioxidant activity for wound healing. Chemical Engineering Journal 2020;392:123775. 10.1016/j.cej.2019.123775.

[83] Nayak BK. Understanding the relevance of sample size calculation. Indian J Ophthalmol 2010;58:469–70. 10.4103/0301-4738.71673.

