## Supplementary material for "Clinical prediction of wound re-epithelisation outcomes in non-severe burn injury using the plasma lipidome": Table S1, Table S2, Figure S1, Figure S2, Figure S3, Figure S4, Figure S5

**Table S1. Annotation of the keys used by the Bruker IVDr Lipoprotein Subclass Analysis (B.I.-LISA™) method for each lipoprotein subclass.** Abbreviations: LDL – low-density lipoprotein; HDL – high-density lipoprotein; VLDL – very low-density lipoprotein; IDL – intermediate-density lipoprotein.

| Key | Class/Subclass | Compound | Concentration unit |
|---|---|---|---|
| TPTG | Total Plasma | Triglycerides | mg/dL |
| TPCH | Total Plasma | Cholesterol | mg/dL |
| TPA1 | Total Plasma | Apolipoprotein-A1 | mg/dL |
| TPA2 | Total Plasma | Apolipoprotein-A2 | mg/dL |
| TPAB | Total Plasma | Apolipoprotein-B100 | mg/dL |
| LDHD | Ratio LDL and HDL Cholesterol | LDL Cholesterol / HDL Cholesterol | -/- |
| ABA1 | Ratio of Apolipoproteins B100 and A1 | Apolipoprotein-B100 / Apolipoprotein-A1 | -/- |
| TBPN | Apolipoprotein-B100 carrying particles | Particle Number | nmol/L |
| VLPN | VLDL | Particle Number | nmol/L |
| IDPN | IDL | Particle Number | nmol/L |
| LDPN | LDL | Particle Number | nmol/L |
| L1PN | LDL-1 | Particle Number | nmol/L |
| L2PN | LDL-2 | Particle Number | nmol/L |
| L3PN | LDL-3 | Particle Number | nmol/L |
| L4PN | LDL-4 | Particle Number | nmol/L |
| L5PN | LDL-5 | Particle Number | nmol/L |
| L6PN | LDL-6 | Particle Number | nmol/L |
| VLTG | VLDL Class | Triglycerides | mg/dL |
| IDTG | IDL Class | Triglycerides | mg/dL |
| LDTG | LDL Class | Triglycerides | mg/dL |
| HDTG | HDL Class | Triglycerides | mg/dL |
| VLCH | VLDL Class | Cholesterol | mg/dL |
| IDCH | IDL Class | Cholesterol | mg/dL |
| LDCH | LDL Class | Cholesterol | mg/dL |
| HDCH | HDL Class | Cholesterol | mg/dL |
| VLFC | VLDL Class | Free Cholesterol | mg/dL |
| IDFC | IDL Class | Free Cholesterol | mg/dL |
| LDFC | LDL Class | Free Cholesterol | mg/dL |
| HDFC | HDL Class | Free Cholesterol | mg/dL |
| VLPL | VLDL Class | Phospholipids | mg/dL |
| IDPL | IDL Class | Phospholipids | mg/dL |
| LDPL | LDL Class | Phospholipids | mg/dL |
| HDPL | HDL Class | Phospholipids | mg/dL |
| HDA1 | HDL Class | Apolipoprotein-A1 | mg/dL |
| HDA2 | HDL Class | Apolipoprotein-A2 | mg/dL |
| VLAB | VLDL Class | Apolipoprotein-B100 | mg/dL |
| IDAB | IDL Class | Apolipoprotein-B100 | mg/dL |
| LDAB | LDL Class | Apolipoprotein-B100 | mg/dL |

| | | | |
|---|---|---|---|
| V1TG | VLDL-1 Subclass | Triglycerides | mg/dL |
| V2TG | VLDL-2 Subclass | Triglycerides | mg/dL |
| V3TG | VLDL-3 Subclass | Triglycerides | mg/dL |
| V4TG | VLDL-4 Subclass | Triglycerides | mg/dL |
| V5TG | VLDL-5 Subclass | Triglycerides | mg/dL |
| V1CH | VLDL-1 Subclass | Cholesterol | mg/dL |
| V2CH | VLDL-2 Subclass | Cholesterol | mg/dL |
| V3CH | VLDL-3 Subclass | Cholesterol | mg/dL |
| V4CH | VLDL-4 Subclass | Cholesterol | mg/dL |
| V5CH | VLDL-5 Subclass | Cholesterol | mg/dL |
| V1FC | VLDL-1 Subclass | Free Cholesterol | mg/dL |
| V2FC | VLDL-2 Subclass | Free Cholesterol | mg/dL |
| V3FC | VLDL-3 Subclass | Free Cholesterol | mg/dL |
| V4FC | VLDL-4 Subclass | Free Cholesterol | mg/dL |
| V5FC | VLDL-5 Subclass | Free Cholesterol | mg/dL |
| V1PL | VLDL-1 Subclass | Phospholipids | mg/dL |
| V2PL | VLDL-2 Subclass | Phospholipids | mg/dL |
| V3PL | VLDL-3 Subclass | Phospholipids | mg/dL |
| V4PL | VLDL-4 Subclass | Phospholipids | mg/dL |
| V5PL | VLDL-5 Subclass | Phospholipids | mg/dL |
| L1TG | LDL-1 Subclass | Triglycerides | mg/dL |
| L2TG | LDL-2 Subclass | Triglycerides | mg/dL |
| L3TG | LDL-3 Subclass | Triglycerides | mg/dL |
| L4TG | LDL-4 Subclass | Triglycerides | mg/dL |
| L5TG | LDL-5 Subclass | Triglycerides | mg/dL |
| L6TG | LDL-6 Subclass | Triglycerides | mg/dL |
| L1CH | LDL-1 Subclass | Cholesterol | mg/dL |
| L2CH | LDL-2 Subclass | Cholesterol | mg/dL |
| L3CH | LDL-3 Subclass | Cholesterol | mg/dL |
| L4CH | LDL-4 Subclass | Cholesterol | mg/dL |
| L5CH | LDL-5 Subclass | Cholesterol | mg/dL |
| L6CH | LDL-6 Subclass | Cholesterol | mg/dL |
| L1FC | LDL-1 Subclass | Free Cholesterol | mg/dL |
| L2FC | LDL-2 Subclass | Free Cholesterol | mg/dL |
| L3FC | LDL-3 Subclass | Free Cholesterol | mg/dL |
| L4FC | LDL-4 Subclass | Free Cholesterol | mg/dL |
| L5FC | LDL-5 Subclass | Free Cholesterol | mg/dL |
| L6FC | LDL-6 Subclass | Free Cholesterol | mg/dL |
| L1PL | LDL-1 Subclass | Phospholipids | mg/dL |
| L2PL | LDL-2 Subclass | Phospholipids | mg/dL |
| L3PL | LDL-3 Subclass | Phospholipids | mg/dL |
| L4PL | LDL-4 Subclass | Phospholipids | mg/dL |
| L5PL | LDL-5 Subclass | Phospholipids | mg/dL |
| L6PL | LDL-6 Subclass | Phospholipids | mg/dL |
| L1AB | LDL-1 Subclass | Apolipoprotein-B100 | mg/dL |
| L2AB | LDL-2 Subclass | Apolipoprotein-B100 | mg/dL |

| | | | |
|---|---|---|---|
| L3AB | LDL-3 Subclass | Apolipoprotein-B100 | mg/dL |
| L4AB | LDL-4 Subclass | Apolipoprotein-B100 | mg/dL |
| L5AB | LDL-5 Subclass | Apolipoprotein-B100 | mg/dL |
| L6AB | LDL-6 Subclass | Apolipoprotein-B100 | mg/dL |
| H1TG | HDL-1 Subclass | Triglycerides | mg/dL |
| H2TG | HDL-2 Subclass | Triglycerides | mg/dL |
| H3TG | HDL-3 Subclass | Triglycerides | mg/dL |
| H4TG | HDL-4 Subclass | Triglycerides | mg/dL |
| H1CH | HDL-1 Subclass | Cholesterol | mg/dL |
| H2CH | HDL-2 Subclass | Cholesterol | mg/dL |
| H3CH | HDL-3 Subclass | Cholesterol | mg/dL |
| H4CH | HDL-4 Subclass | Cholesterol | mg/dL |
| H1FC | HDL-1 Subclass | Free Cholesterol | mg/dL |
| H2FC | HDL-2 Subclass | Free Cholesterol | mg/dL |
| H3FC | HDL-3 Subclass | Free Cholesterol | mg/dL |
| H4FC | HDL-4 Subclass | Free Cholesterol | mg/dL |
| H1PL | HDL-1 Subclass | Phospholipids | mg/dL |
| H2PL | HDL-2 Subclass | Phospholipids | mg/dL |
| H3PL | HDL-3 Subclass | Phospholipids | mg/dL |
| H4PL | HDL-4 Subclass | Phospholipids | mg/dL |
| H1A1 | HDL-1 Subclass | Apolipoprotein-A1 | mg/dL |
| H2A1 | HDL-2 Subclass | Apolipoprotein-A1 | mg/dL |
| H3A1 | HDL-3 Subclass | Apolipoprotein-A1 | mg/dL |
| H4A1 | HDL-4 Subclass | Apolipoprotein-A1 | mg/dL |
| H1A2 | HDL-1 Subclass | Apolipoprotein-A2 | mg/dL |
| H2A2 | HDL-2 Subclass | Apolipoprotein-A2 | mg/dL |
| H3A2 | HDL-3 Subclass | Apolipoprotein-A2 | mg/dL |
| H4A2 | HDL-4 Subclass | Apolipoprotein-A2 | mg/dL |

**Table S2. List of de-identified assigned patient numbers and plasma successfully analysed by LC-QQQ-MS and NMR spectroscopy at each timepoint collection.**

| | Burns admission | Surgery | Two days post-surgery | Two weeks post-surgery | Six weeks post-surgery |
|---|---|---|---|---|---|
| *Patient 1* | ✓ | ✓ | ✓ | ✓ | ✓ |
| *Patient 2* | ✓ | ✓ | | ✓ | ✓ |
| *Patient 3* | ✓ | ✓ | | ✓ | |
| *Patient 4* | ✓ | | ✓ | | ✓ |
| *Patient 5* | ✓ | | ✓ | ✓ | ✓ |
| *Patient 6* | ✓ | | ✓ | ✓ | ✓ |
| *Patient 7* | ✓ | ✓ | | ✓ | ✓ |
| *Patient 8* | ✓ | ✓ | ✓ | | |
| *Patient 9* | ✓ | ✓ | ✓ | ✓ | ✓ |
| *Patient 10* | | ✓ | | | |
| *Patient 11* | | | ✓ | | ✓ |
| *Patient 12* | ✓ | | ✓ | | ✓ |
| *Patient 13* | ✓ | | ✓ | | |
| *Patient 14* | ✓ | | | ✓ | ✓ |
| *Patient 15* | ✓ | ✓ | | ✓ | ✓ |
| *Patient 16* | ✓ | ✓ | | | ✓ |
| *Patient 17* | ✓ | | | ✓ | ✓ |
| *Patient 18* | ✓ | | ✓ | | ✓ |
| *Patient 19* | ✓ | | ✓ | ✓ | ✓ |
| *Patient 20* | ✓ | ✓ | | | ✓ |
| ***Total*** | **18** | **10** | **11** | **11** | **16** |

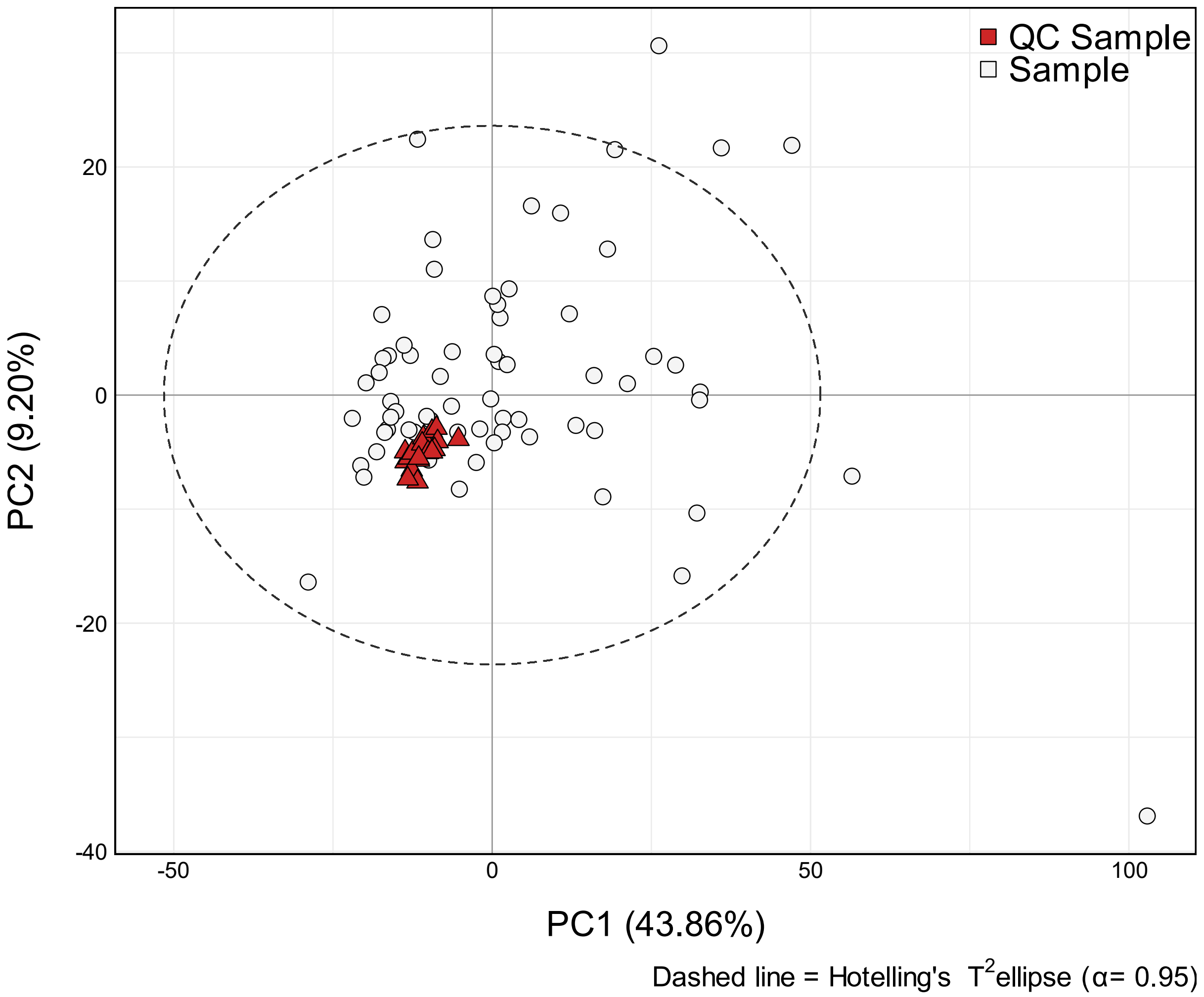


**Figure S1. Principal component analysis (PCA) of non-severe burn plasma samples at all timepoints and intermittently analysed plasma quality control (QC) samples.** Scores plot of cohort (n = 66; white circle) and QC samples (n=18; red triangle).

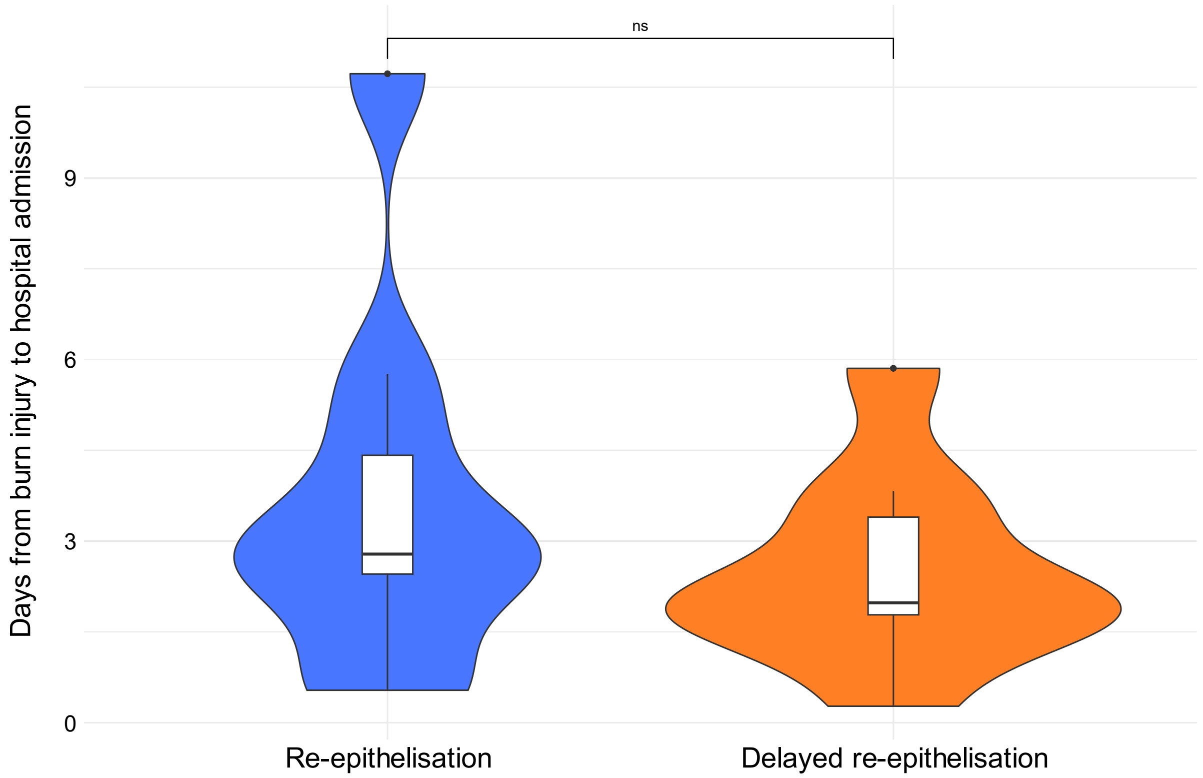


**Figure S2. Combined violin and box and whisker plot of days from burn injury to hospital admission in re-epithelisation groups at two weeks post-surgery for the non-severe burn cohort.** Re-epithelialized (n=9; coloured blue) and delayed re-epithelisation (n=9; coloured orange) at admission were compared using Mann-Whitney U univariate analysis. Significance: ns – not significant (p-value > 0.05).

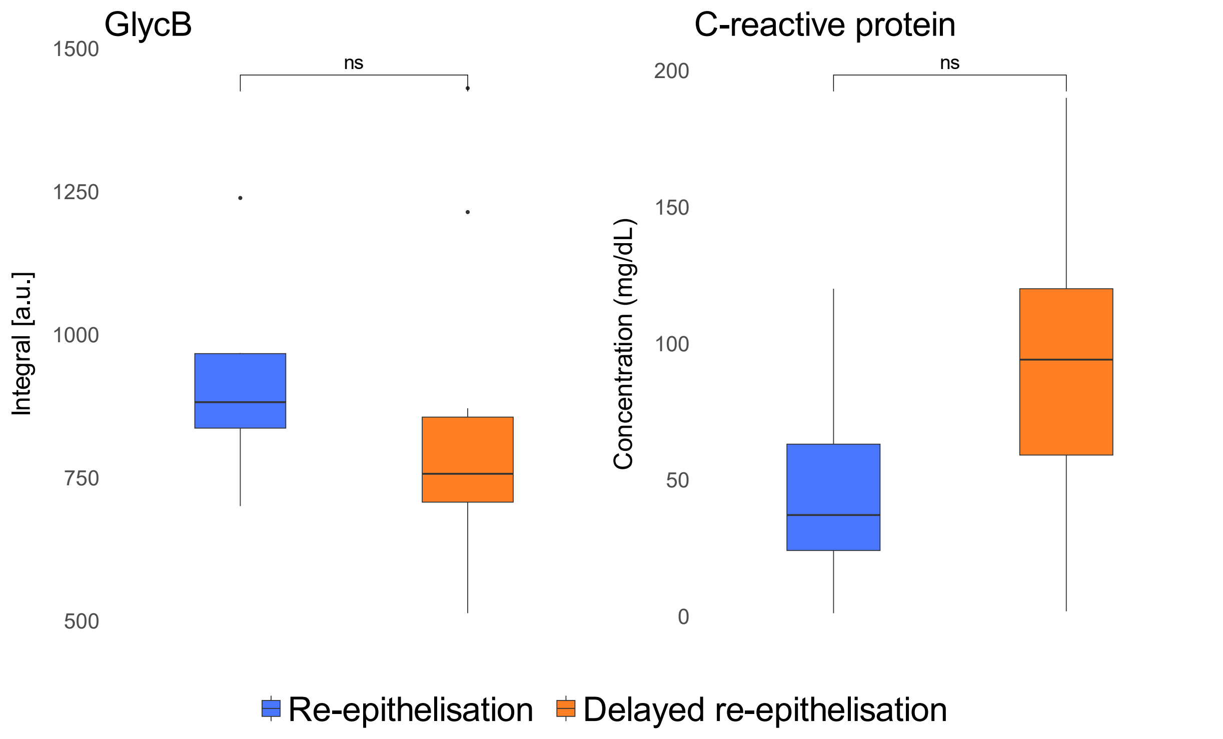


**Figure S3. Box and whisker plots of GlycB and C-reactive protein plasma levels in the two epithelisation groups at admission for the non-severe burn cohort.** Re-epithelialized (n=9; coloured blue) and delayed re-epithelisation (n=9; coloured orange) at admission were compared using Mann-Whitney U univariate analysis. Significance: ns – not significant (p-value > 0.05).

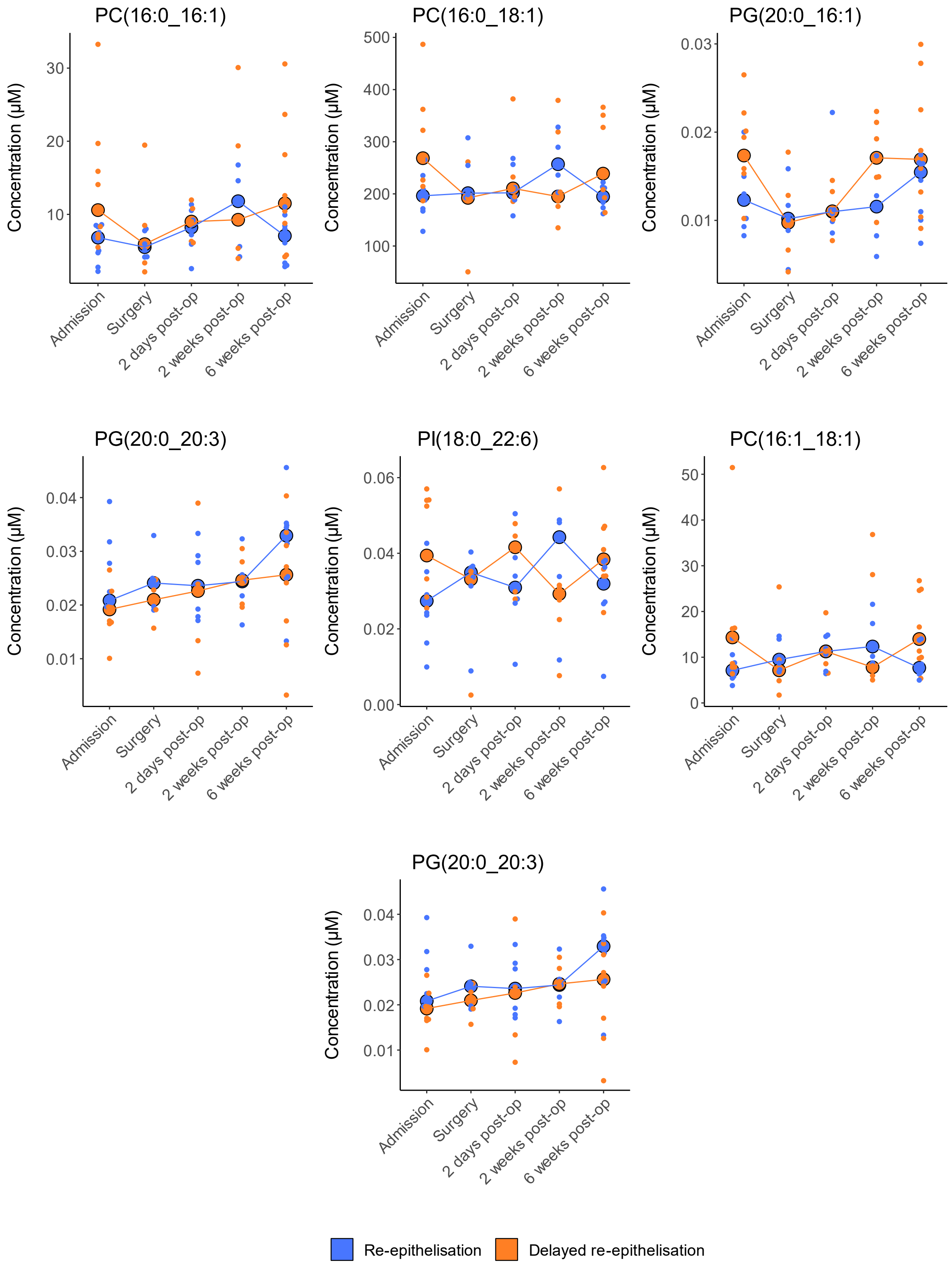


**Figure S4. Trajectory dot plots of 7 significant and main separation driver lipid species from the eruption plot and OPLS-DA modelling for both wound outcome groups over time**. Plots display lipid concentrations of each patient with median trendline of both RE (n=10; coloured blue) and DRE (n=10; coloured orange) at burns admission, surgery, two days, two weeks and six weeks-post surgery.

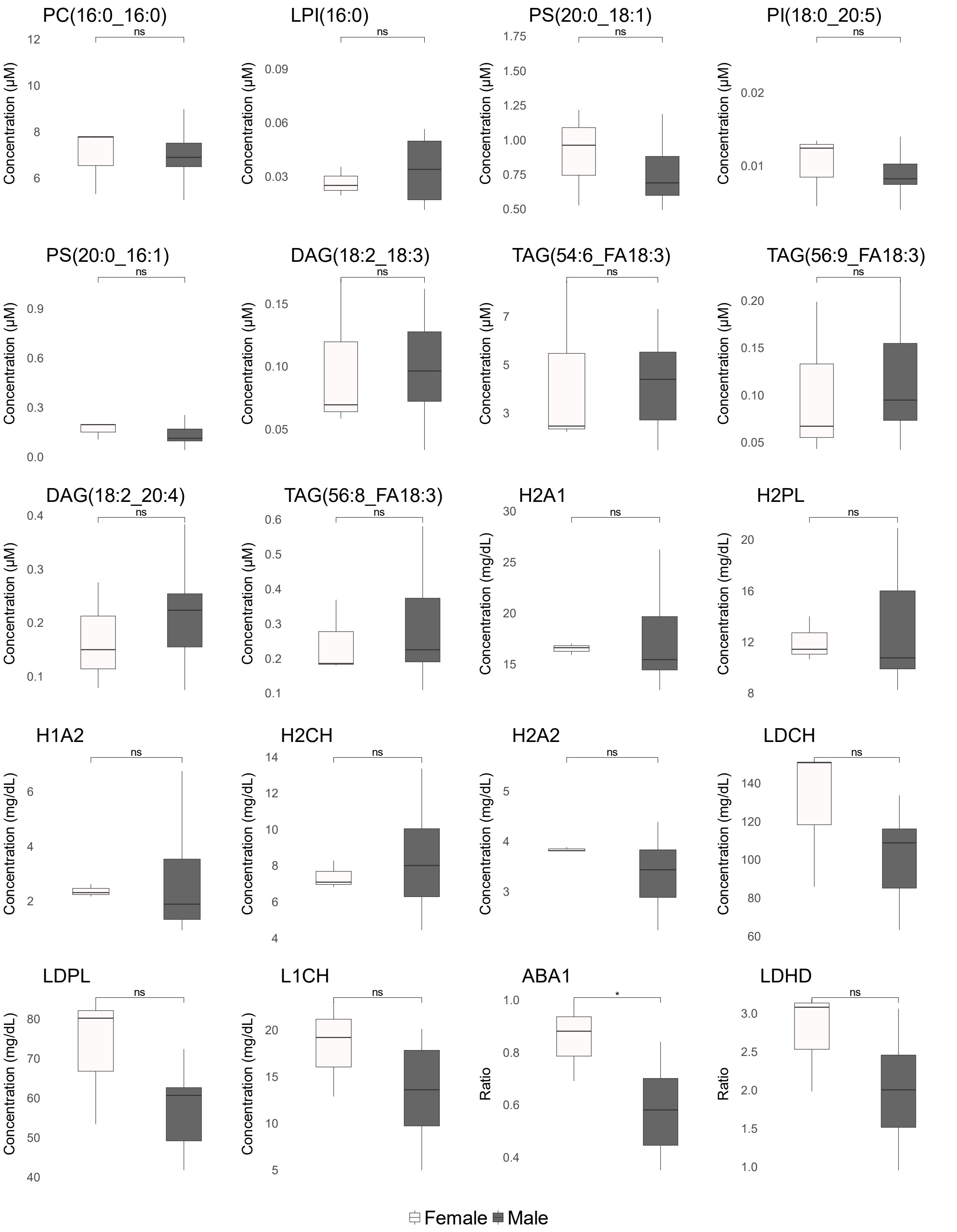


**Figure S5. Box and whisker plots of 20 key lipids and lipoproteins driving re-epithelisation group separation to assess any influence of sex.** Differences in female (n = 3, coloured white) and male (n = 17, coloured grey) sexes for key lipids were analysed by Mann-Whitney U at burns admission. Significance: ns – not significant (p-value > 0.05), '*' = p-value < 0.05.
